# Mapping Higher-Order Topology in OCD Brain Networks with Hodge Laplacian

**DOI:** 10.64898/2026.03.04.709586

**Authors:** Hanyang Ruan, Moo K. Chung, Willem B. Bruin, Nadža Džinalija, Yoshinari Abe, Pino Alonso, Alan Anticevic, Srinivas Balachander, Marcelo C. Batistuzzo, Francesco Benedetti, Sara Bertolín, Silvia Brem, Youngsun Cho, Federica Colombo, Beatriz Couto, Goi Khia Eng, Sónia Ferreira, Jamie D. Feusner, Rachael G. Grazioplene, Patricia Gruner, Kristen Hagen, Bjarne Hansen, Yoshiyuki Hirano, Marcelo Q. Hoexter, Jonathan Ipser, Fern Jaspers-Fayer, Minah Kim, Jun Soo Kwon, Luisa Lazaro, Chiang-Shan R. Li, Christine Lochner, Rachel Marsh, Ignacio Martínez-Zalacaín, Jose M. Menchón, Pedro S. Moreira, Pedro Morgado, Emma Muñoz, Akiko Nakagawa, Janardhanan C. Narayanaswamy, Erika L. Nurmi, Joseph O’Neill, Jose C. Pariente, John C. Piacentini, Maria Picó-Pérez, Fabrizio Piras, Federica Piras, Christopher Pittenger, Janardhan Y. C. Reddy, Daniela Rodriguez-Manrique, Yuki Sakai, Joao R. Sato, Eiji Shimizu, Venkataram Shivakumar, Helen B. Simpson, Carles Soriano-Mas, Nuno Sousa, Emily R. Stern, Evelyn Stewart, Philip R. Szeszko, Sophia I. Thomopoulos, Anders L. Thorsen, Benedetta Vai, Anouk van der Straten, Ysbrand D. van der Werf, Wieke van Leeuwen, Hein van Marle, Guido van Wingen, Daniela Vecchio, Ganesan Venkatasubramanian, Chris Vriend, Susanne Walitza, Zhen Wang, Tokiko Yoshida, Je-Yeon Yun, Qing Zhao, ENIGMA OCD Working Group, Paul M. Thompson, Dan J. Stein, Odile A. van den Heuvel, Kathrin Koch

## Abstract

Brain disorders are increasingly understood as disorders of distributed brain circuits, yet functional connectivity (FC), the dominant framework for mapping them, treats the brain as a collection of pairwise relationships between regions and cannot represent pathology distributed across coordinated sets of connections. We introduce a Hodge-Laplacian topological framework that localizes higher-order “loop” (1-cycle) organization within functional connectome, maps each loop to specific edges and networks, and yields a subject-level measure of loop expression. Applied to resting-state fMRI from the ENIGMA-OCD consortium (1,024 patients and 1,028 controls across 28 sites), the framework identified 93 loop-level abnormalities in obsessive–compulsive disorder (OCD), concentrated in frontoparietal and somatomotor systems. The edges forming these loops largely showed no significant differences between groups, indicating that the abnormalities were invisible to conventional FC analysis. The frontoparietal and somatomotor loop clusters recurred across the clinical subgroups, suggesting convergence on a shared higher-order phenotype. Robustness analyses showed the loop signal reflected higher-order organization rather than an artifact of individual edges, the network backbone, or any single site. These results indicate that coordinated, multi-edge pathology exists and can be localized even when pairwise analyses fail to detect it, positioning higher-order topology as a generalizable axis for mapping circuit pathology across psychiatric and neurological disorders.

## Introduction

Psychiatric and neurological disorders are increasingly modelled as disorders of distributed brain circuits, instead of collections of isolated abnormal brain regions (1, 2). Resting-state functional magnetic resonance imaging (rs-fMRI), as one of the principal tools for investigating brain network function, has provided evidence for these altered circuit connectivities in many psychiatric disorders(3–5). However, conventional rs-fMRI analyses have focused on pairwise functional connectivity (FC), assuming that network dysfunction can be decomposed by treating the brain as a collection of interactions between pairs of regions. This decomposition is powerful, but cannot represent pathology arising from the simultaneous organization of multiple connections. This matters because the leading neurobiological hypotheses of psychiatric disorders, for instance the cortico-striatal-thalamic-cortical (CSTC) circuit in obsessive-compulsive disorder (OCD) and the triple-network dysfunction in schizophrenia, are inherently defined by loops rather than isolated connections. An edge-centric description can therefore potentially miss multi-regional loop-like alterations even when individual pairwise FC changes are not detectable (6–9).

Capturing this organization requires extending the analysis from pairs of regions to loop-like structures formed by three or more regions, and topological data analysis (TDA) offers a principled way to achieve this. TDA is an emerging mathematical framework grounded in algebraic topology for probing the shape and structure of data (10). It is especially suitable for data that are intrinsically complex, multi-dimensional and non-linear, and has been applied across domains from molecular structure, genomics to medical imaging (11–14) By modeling brain networks as simplicial complexes rather than simple graphs based on pairwise FC, TDA tracks higher-order structure via persistent homology (PH) (15–17). Several studies have applied PH to fMRI, revealing novel biomarkers in various psychiatric disorders (18–21). However, while PH can quantify the presence of abnormal loops (e.g., via Betti numbers), it cannot locate which regions or edges form them—a spatial ambiguity that hinders clinical interpretation.

Recently, the Hodge Laplacian has been proposed to overcome this limitation. Unlike conventional PH approaches, it enables spatial localization of topological features and has been used to detect and locate cyclic structures in domains ranging from cell-development trajectories to brain networks (22–24). It assesses physiologically inherent higher-order structures - 1-cycles, or loops - and localizes them with regard to canonical brain networks. A 1-cycle encodes a closed, multi-edge route, as a minimal representation for higher-order interactions across distributed regions that can be potentially linked to feedforward and feedback processes (25, 26). The Hodge Laplacian framework makes it possible to test disease effects directly on specific loops rather than single edges. This cycle-centric view is aligned with the broader idea that cyclic interactions form the foundation of brain dynamics regarding complex cognitive processes (27, 28), and offers an organizational axis along which psychopathology may manifest (29, 30). An overview of the conventional TDA and the Hodge Laplacian on a graph is presented in Figure 1A.

**Fig. 1.**
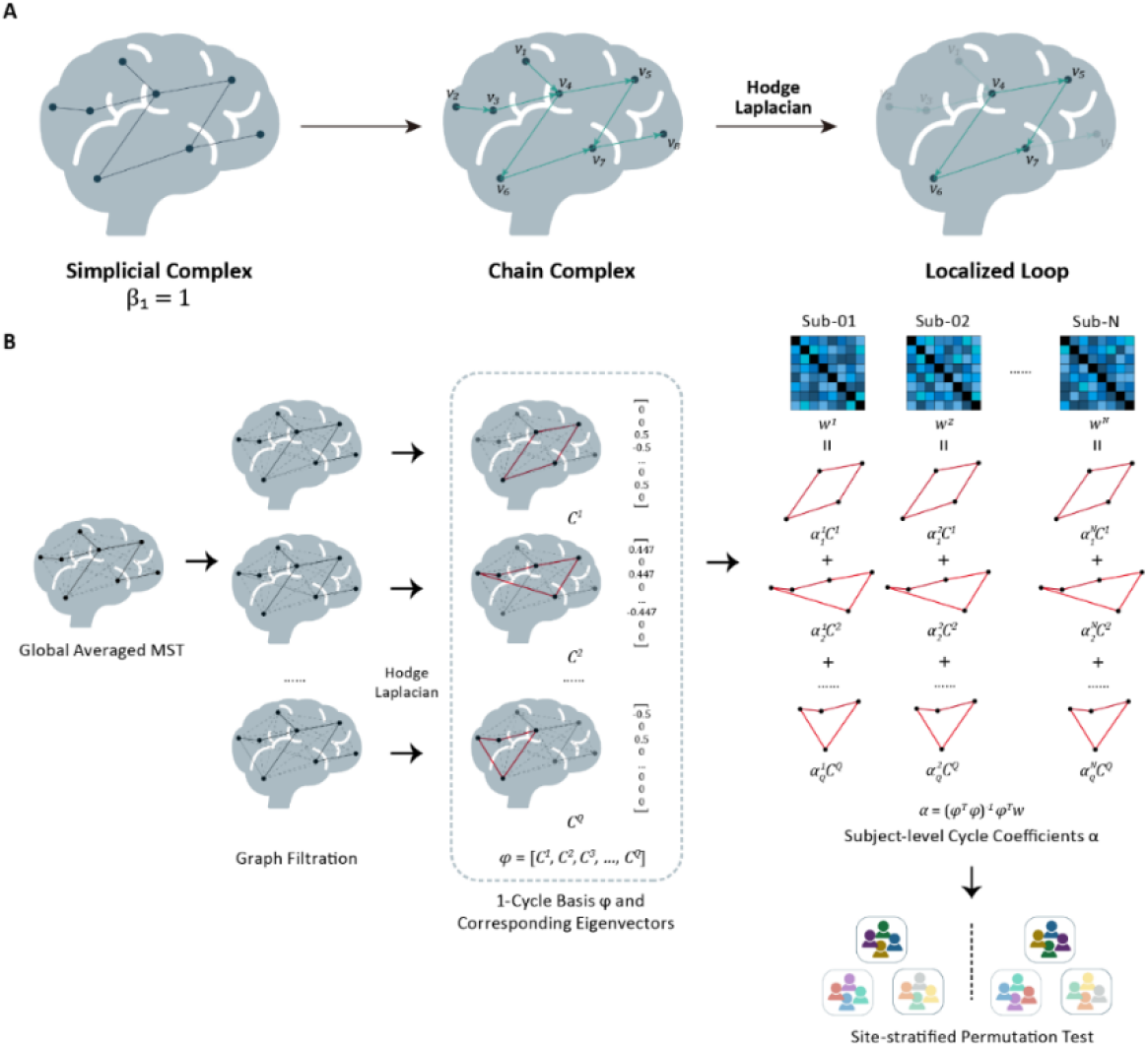
Overview of the Hodge Laplacian Topological Analysis. **A**. Conceptual Illustration. Traditional topological data analysis (left) can detect whether a loop exists in a topological space (e.g., Betti number *β*_1_ = 1, indicating existence of 1 loop), but it does not identify where the loop is located or which edges contribute to it. The Hodge Laplacian framework uses the chain complex (middle), where edges are assigned with directions and organized into boundary operators. Decomposing edge flows using the Hodge Laplacian separates them into gradient, curl, and harmonic components. The harmonic component captures non-trivial cycles that are not boundaries of higher-dimensional structures. By isolating this component, we can localize the specific edges that form the loop (right), enabling spatial interpretation within the brain network. **B**. Computation of the Hodge Laplacian on resting state brain networks. First, we construct a group-level network skeleton using the maximum spanning tree (MST) to ensure connectivity while minimizing redundancy. From this skeleton, we derive a common 1-cycle basis representing fundamental loops in the group-averaged topology. Individual brain networks are then projected onto the 1-cycle basis. The resulting cycle coefficients represent the expression of each loop in each subject. These subject-specific cycle coefficients are used as features for downstream statistical analysis (e.g., group comparisons or correlations with clinical measures in OCD).

In this study, we introduce a Hodge Laplacian-based TDA framework for functional connectomes that localizes higher-order loop pathology and produces a per-subject cycle measure. We validate it on a large-scale resting-state fMRI dataset from the ENIGMA-OCD working group, the largest multi-site OCD cohort assembled to date. OCD is a disabling psychiatric condition characterized by intrusive obsessions and repetitive compulsions (31). Conventional neuroimaging has proposed circuitry hypotheses such as CSTC, frontal-limbic and sensorimotor circuits, but these were all based on pairwise FC results (29, 32, 33), and the reported group-level edge effects were small (34). We hypothesize that OCD involves disruptions of higher-order network organization, particularly within large-scale networks related to cognitive control and somatomotor systems, can be identified as localized 1-cycle abnormalities, but remain largely invisible to conventional FC analysis. Broadly, this framework will also offer a generalizable axis for detecting higher-order circuit pathology across psychiatric and neurological disorders.

## Results

### Higher-order loops reveal OCD network pathology invisible to pairwise connectivity

To investigate higher-order topological abnormalities in OCD brain networks, we analyzed resting-state functional MRI (rs-fMRI) data from 2,052 participants (1,024 patients and 1,028 healthy controls) from the ENIGMA-OCD working group (Supplementary Table S1). Using a Hodge Laplacian framework (Fig. 1B), we constructed a common 1-cycle basis from the group-averaged FC matrix, projected each subject’s FC onto it, and tested the resulting 1-cycle coefficients for group differences with a site-stratified Freedman-Lane permutation integrating general linear model (GLM) to adjust age, sex, head motion, and scan site effects (35).

We identified 93 significant discriminating 1-cycles in individuals with OCD compared to controls (family-wise error rate (FWER) corrected *p*<0.0001; Cohen’s *d* = 0.223-0.283, partial R^2^ = 0.012-0.019; null-distribution in Supplementary Fig. S3). Agglomerative clustering of their functional profiles revealed five clusters spanning the frontoparietal network (FPN), visual network (Vis)/dorsal attention network (DAN), somatomotor network (SMN), default mode network (DMN), and SMN/salience network (SN), with most cycles in the intra-FPN and intra-SMN clusters (Fig. 2A, B). Cluster-level properties and the most discriminating cycle per family are shown in Fig. 2C.

**Fig. 2.**
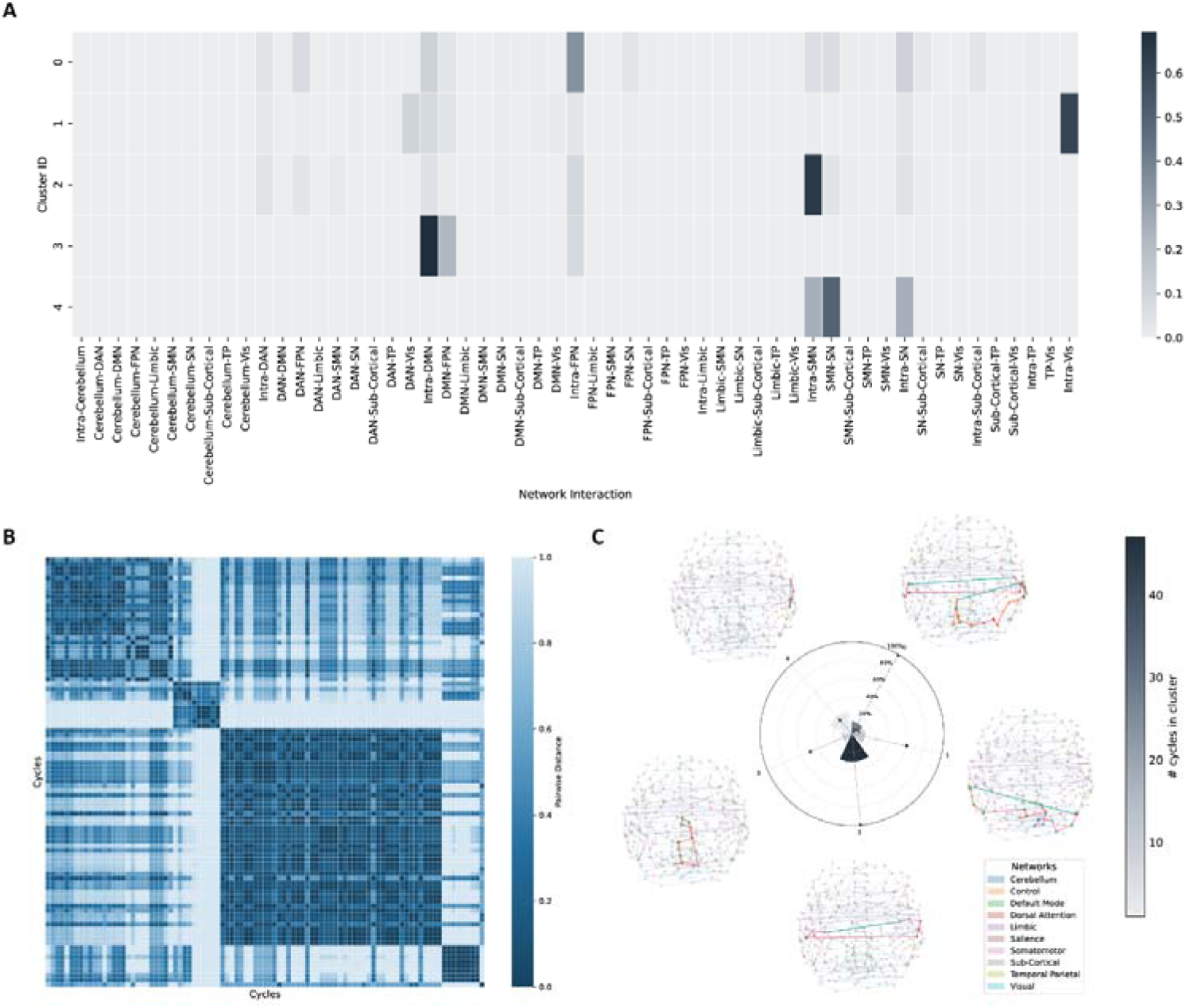
1-cycle abnormalities in main analysis. **A**. Agglomerative clustering results according to functional profiles of 1-cycles. **B**. Pairwise cosine distances between cycles. 5 distinct clusters are shown in the heatmap. Cosine distance was computed between normalized 55-dimensional network-interaction profiles, smaller values indicate similar network composition. **C**. Radial bar plot summarizing the following three cluster-level properties: Wedge height indicates the mean fraction of edges per 1-cycle that show significant edge-wise FC differences between OCD and controls, averaged across 1-cycles within each cluster. Bar color illustrates the number of significant 1-cycles in the cluster. The dot marks the cluster’s normalized mean 1-cycle length (mean number of edges per 1-cycle, normalized to the maximum across clusters). Visualizations of the most discriminating 1-cycles in each cluster are displayed around the plot. Global averaged maximum spanning tree structure is shown in the background. Different colors of nodes indicate different brain networks. Highlighted edges demonstrate the higher-order cycle structure, with green color indicating a significant difference in the functional connectivity strength of this edge between OCD and healthy controls. DAN: dorsal attention network, DMN: default mode network, FPN: frontoparietal network (labeled “Control” in the atlas), OCD: obsessive-compulsive disorder, SMN: somatomotor network, SN: salience network, TP: temporal parietal network, Vis: visual network.

Critically, these alterations were largely invisible to conventional edge-wise FC: most edges composing the abnormal cycles showed no significant pairwise group difference in a linear mixed-effects (LME) model (non-green edges, Fig. 2C), confirming that the 1-cycles capture higher-order, multi-nodal disruptions. Specifically, individuals with OCD exhibited weaker FCs in these edges, indicating weaker integration in these network-specific cycles that mostly spanned both hemispheres.

### Loop pathology converges on the same networks across clinical subgroups

To relate these alterations to clinical features, we applied the same framework to subgroups stratified by age, medication status, severity, and age of onset, each compared against HC (Table 1). We projected each significant subgroup cycle onto the five clusters from main-analysis by cosine similarity of their 55-dimensional interaction profiles (independent per-subgroup clustering in Supplementary Figures).

**Table 1.** Summary of permutation results regarding OCD vs HC across the main and subgroup analyses. Global *p* denotes the FWER-corrected permutation *p* value for the group effect. “Cycles (N)” indicates the number of significant 1-cycles. Effect sizes (Cohen’s *d*, partial R^2^) are reported as ranges across significant 1-cycles.

| | OCD (N) | HC (N) | Global $p$ | Cohen's $d$ | Partial $R^2$ | Cycles (N) |
| --- | --- | --- | --- | --- | --- | --- |
| Pediatric | 103 | 101 | 0.839 | / | / | / |
| Adult | 903 | 914 | 0.0001 | 0.237-0.304 | 0.013-0.021 | 63 |
| Unmedicated | 356 | 420 | 0.0139 | 0.381 | 0.034 | 1 |
| Medicated | 456 | 683 | 0.0001 | 0.307-0.474 | 0.020-0.047 | 179 |
| Low-severity | 501 | 880 | 0.0056 | 0.285-0.309 | 0.017-0.021 | 6 |
| High-severity | 414 | 762 | 0.0001 | 0.306-0.410 | 0.020-0.036 | 55 |
| Early-onset | 305 | 567 | 0.01 | 0.369-0.387 | 0.028-0.031 | 5 |
| Adult-onset | 388 | 663 | 0.0005 | 0.334-0.390 | 0.023-0.031 | 12 |
| Main | 1024 | 1028 | 0.0001 | 0.223-0.283 | 0.012-0.019 | 93 |

Across subgroups with detectable effects, significant loops projected onto the same intra-FPN and intra-SMN clusters, indicating convergence on a common higher-order phenotype rather than distinct per-subgroup topologies (Fig. 3). Loop counts scaled with sample size while per-cycle effect sizes were comparable, so differences in counts reflect detection power rather than the magnitude of loop-level pathology: effects were strongest in the adult, medicated, and high-severity subgroups and absent in the pediatric sample (Table 1). In an exploratory test, the intra-FPN loop-expression score was positively associated with Y-BOCS severity after adjustment for age, sex, head motion, and site (*t* = 3.19, partial *r* = 0.102, false discovery rate (FDR)-corrected *p* = 0.0074; Supplementary Table S3).

**Fig. 3.**
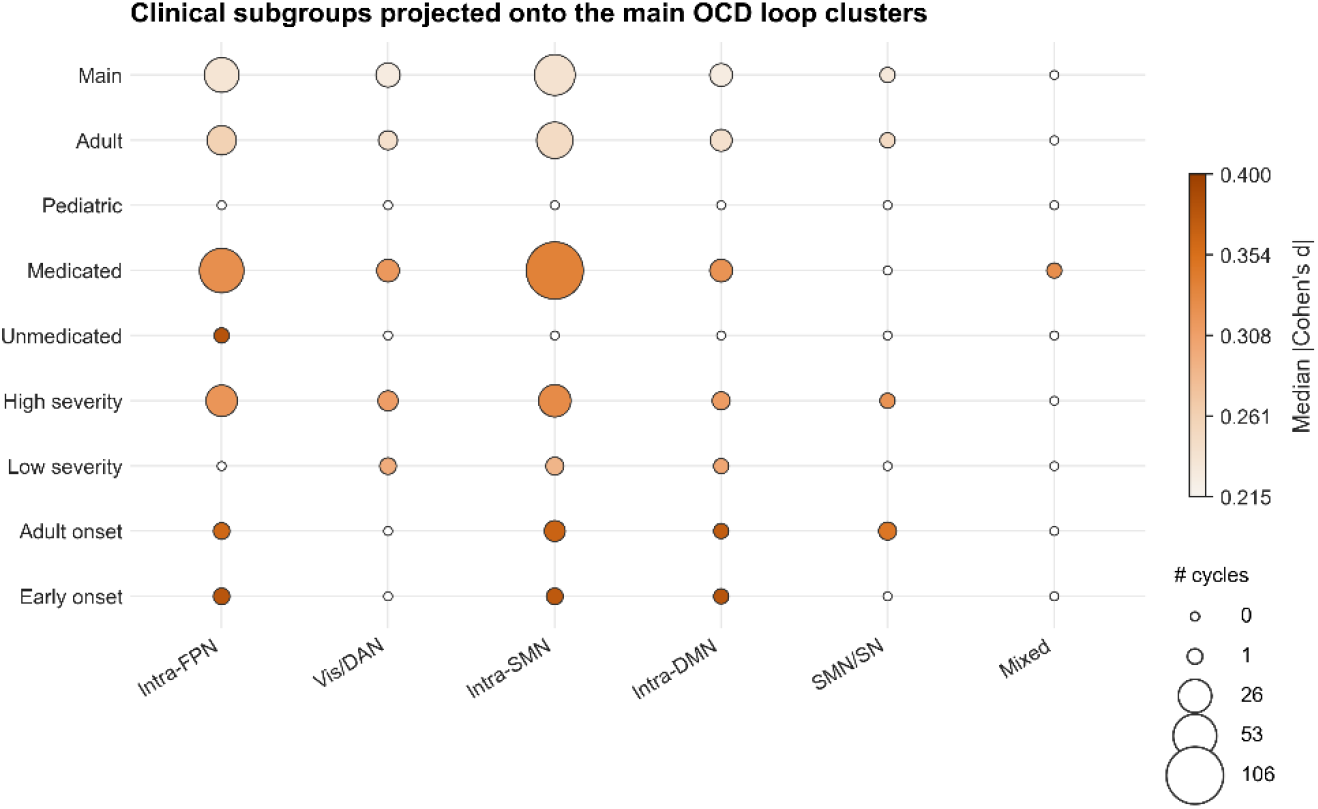
Clinical subgroup projection of OCD-related higher-order loop abnormalities. Significant 1-cycles from each clinical subgroup analysis mapped onto the 5 clusters identified in the main analysis. Bubble size denotes the number of significant 1-cycles assigned to each main-analysis family, and bubble color denotes the median absolute Cohen’s *d*. Small white circles denote zero significant cycles. Adult, medicated, and high-severity OCD subgroups showed broader involvement of the main OCD loop clusters, particularly intra-SMN and intra-FPN cycles, whereas pediatric, unmedicated, and low-severity subgroups showed limited or no detectable loop-level abnormalities.

### The loop-level signature is specific, independent, and robust across sites

Three additional analyses were performed to establish specificity and robustness. First, to test whether the workflow could generate spurious topological differences in the absence of disease, we repeated the complete workflow in a null split-half analysis of healthy controls. In this case any detected significant 1-cycle difference would indicate sampling- or pipeline-induced topology rather than disease effect. No split-half difference was detected (*p* = 0.429). Second, because of the nature of graph filtration (SI methods), the maximum spanning tree (MST) of the graph is used to define the common 1-cycle basis: each off-tree edge closes one such cycle. We therefore separated each cycle coefficient into backbone-related and loop-closing contributions to test whether the group effect was driven by the spanning-tree itself or by the edge that completes the cycle. We showed that all 93 cycles differed on the off-tree component but only 41 on the tree component, confirming that significance reflected genuine loop closure by non-MST edges rather than the spanning-tree backbone (Supplementary Table S2). Third, under a leave-one-site-out (LOSO) scheme, cluster-level effects were moderate (Cohen’s *d* = 0.23–0.51) and highly stable: removing any site shifted each estimate only marginally and no LOSO interval crossed zero (intra-FPN *d* = 0.51 [0.48, 0.52]; intra-DMN 0.43 [0.35, 0.44]; intra-SMN 0.42 [0.38, 0.44]; Vis/DAN 0.41 [0.39, 0.44]; SMN/SN 0.23 [0.14, 0.24]; Fig. 4). Within-site effects were positive at the large majority of sites for all five clusters (Supplementary Fig. S11), confirming a broadly shared signature rather than one driven by a few centers.

**Fig 4.**
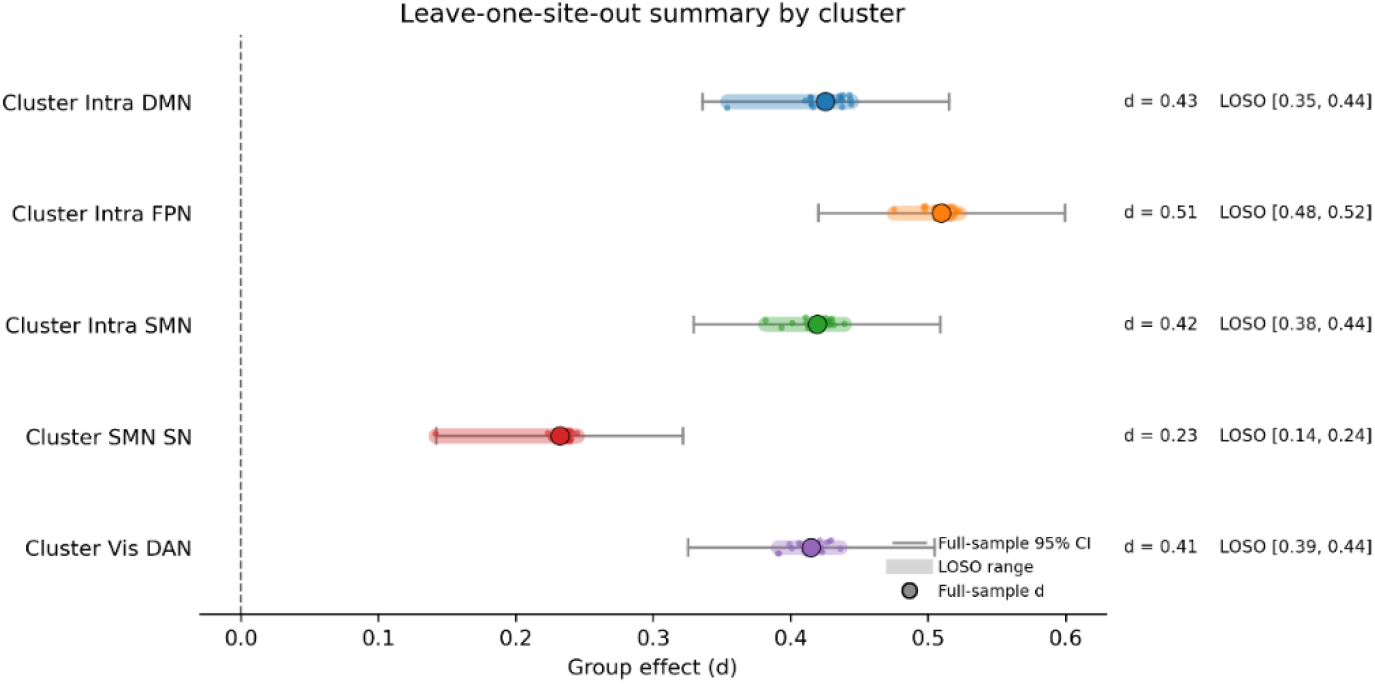
Forest plot of leave-one-site-out stability of cluster-level effects. Cluster-level OCD–HC effects remained positive and highly stable after omitting any single acquisition site. Large dots denote full-sample Cohen’s *d*, grey bars denote full-sample 95% confidence intervals, small dots denote leave-one-site-out estimates, and shaded bands denote the corresponding leave-one-site-out range.

## Discussion

In this study we introduced a Hodge-Laplacian-based TDA framework for functional brain networks that localizes abnormal 1-cycles, and supports group-level inference of higher-order disruptions. In a multi-site resting-state mega-analysis of the ENIGMA-OCD cohort, results showed distinct groups of altered cycles in OCD spanning FPN, Vis/DAN, SMN, DMN, and SMN/SN networks, and the same predominantly FPN- and SMN-centered clusters recurred across adult, medicated, and high-severity subgroups. The edges constituting these discriminating 1-cycles largely lacked significant pairwise group differences, so the abnormalities were visible only at the higher-order level and not in conventional FC analysis. Control analyses further showed that these higher-order effects are reliable, unique to loop structure and robust across multi-sites. Together, these findings extend current models of OCD pathology by highlighting disruptions in higher-order loop organization, in addition to isolated pairwise connections.

The most discriminating 1-cycles were primarily composed of edges without significant between-group differences in edge-wise LME analysis, indicating that OCD-related abnormalities are not only reflected in individual connections but also how edges cohere into coordinated multi-node routes. By elevating inference from edges to loops, our results suggest that OCD pathophysiology may involve a disruption in higher-order coordination among large-scale systems. This is consistent with the long-standing view that OCD is a disorder with distributed abnormal circuit dysfunction (33, 36, 37), while adding a new, localizable descriptor of higher-order cyclic structures that is largely invisible to standard FC analyses.

The disrupted 1-cycles were identified predominantly in the FPN, DMN, SMN, and visual networks, which are known to be closely related to impaired cognitive control and somatomotor function in OCD, as suggested by prior resting state studies (5, 38–41). FPN-DMN interactions support the flexible alternation between externally oriented executive control and internally generated thoughts, with these abnormalities potentially causing impaired control over internally generated signals (34, 42, 43). SMN and visual regions are related to the generation and monitoring of motor behavior and sensory information (33, 44). Clinical studies indicated that altered motor control and sensory phenomena are experienced by a large proportion of OCD patients, which contributed to repetitive compulsive behaviors like ordering, counting or arranging (38, 45).

Our results therefore extend the current understanding of OCD pathophysiology by indicating higher-order loop abnormalities in networks involved in these cognitive processes and functions, even when individual edges show only subtle changes. We hypothesize a hierarchical organization of OCD-related brain network alterations, in which distributed, multi-edge topological disruptions are superimposed on more localized edge-wise differences reported previously. In addition, subgroup analyses showed a convergent pattern of predominant FPN- and SMN-related higher-order disruptions, typically in adult, medicated and high-severity patients. This mirrors prior ENIGMA-OCD resting-state findings at the edge level, where ROI-to-ROI differences were most evident in adult, medicated and high-severity subgroups (34). Such clinical burden related effects also appeared in white matter and brain volume abnormalities in OCD patients (46–48). An exploratory dimensional analysis showed that greater intra-FPN loop expression was associated with higher Yale-Brown Obsessive Compulsive Scale (Y-BOCS) severity in OCD patients. It suggests that the FPN-centered loop phenotype may be clinically relevant rather than only distinguishing cases from controls. Together, these results suggest that both pairwise and higher-order abnormalities may reflect a more trait-like re-organization of the brain that becomes prominent with greater cumulative symptom burden, and/or treatment exposure.

A central question for any higher-order method is whether the detected structure is genuine. Although evidence from TDA and hypergraph analysis has showed that the brain network is a complex system beyond simple pairwise interactions (6, 18), topological features of dimension two and above cannot be stably estimated, especially across multiple subjects (49). Our control analyses indicate that the 1-cycle effects reflect true signal rather than artifacts: a split-half analysis in healthy controls produced no spurious group topology; decomposing each cycle showed that the effects depended on the cycle-closing (off-tree) edges rather than the spanning-tree backbone, confirming a genuinely higher-order origin; and within-site and LOSO analyses showed that no single site drove the result. Together these results showed the robustness of the Hodge Laplacian framework, which can be generalized to other psychiatric conditions and large dataset, provide reliable estimation of higher-order 1-cycle structures in functional brain networks, and potentially extend the current understanding of disease models.

Several limitations should be considered. First, given the cross-sectional nature of the ENIGMA dataset, we cannot determine the direction of causality. As in previous ENIGMA-OCD analyses, we found significant topological disruptions especially in medicated and adult subgroups. It is difficult to distinguish whether stronger 1-cycle abnormalities reflect treatment exposure, greater severity, longer duration, or a trait marker for OCD pathology. Second, to ensure statistical comparability across thousands of subjects, we projected individual data onto a group-level 1-cycle basis derived from the group-averaged network. While this approach robustly identifies shared pathological backbones, it potentially neglects some individualized topological features, while also restraining the interpretation inside this specific dataset. Third, our analysis was performed on samples from existing studies worldwide, using different scanners and without harmonized data collection protocols. Although we have used a site-stratified permutation scheme together with correction of site-effects in the GLM model to minimize the heterogeneity of the multi-site data, and the LOSO results indicate the effects are not site-driven, the presence of additional (e.g. scanner-related) site effects cannot fully be excluded. Finally, we restricted our analysis to 1-dimensional harmonic cycles. While it’s statistically robust, it leaves higher-dimensional organization unexamined.

In conclusion, the Hodge Laplacian framework allows us to unveil higher-order functional abnormalities of OCD. We identified a set of disrupted, weakened higher-order functional loops, spanning frontoparietal, default mode, somatomotor, and visual networks. Importantly, these higher-order loop abnormalities cannot be explained by pairwise comparisons and suggest a circuit-based brain architecture in which pathology manifests not only as global pairwise dysconnectivity, but as a hierarchical disruption of distributed higher-order topological integration. By moving beyond the ‘edge-centric’ view to a ‘cycle-centric’ perspective, our framework provides a robust, mathematically rigorous, and generalizable tool to map the disrupted circuitries that might be associated with psychiatric symptomatology. As psychiatric disorders are increasingly understood as disorders of distributed circuit dynamics rather than isolated regional lesions (50–52), we expect this approach to be broadly useful for probing the still largely unexplored higher-order topology of psychopathology, in OCD and beyond.

## Materials and Methods

### Study samples

We analyzed data provided by the ENIGMA-OCD working group (https://enigma.ini.usc.edu/ongoing/enigma-ocd-working-group/). 1,024 OCD patients and 1,028 healthy controls from 28 sites, in total 2,052 subjects, were finally included in the study (see SI Methods for detailed inclusion/exclusion criteria). The diagnosis of OCD was based on DSM-IV(-TR) or DSM-5 criteria, the Y-BOCS and the Children’s Y-BOCS were used for assessing symptom severity. All healthy participants were free of past and present diagnosis of psychiatric disorders as well as free of any psychotropic medication use at the time of inclusion. All studies were approved by the local institutional review board and participants provided written informed consent.

### Image acquisition and processing

Structural T1-weighted and resting-state functional MRI data were acquired at 1.5 or 3 tesla and preprocessed locally at each site. rs-fMRI data were obtained for 4–12□min with a repetition time ranging between 700 and 3500□ms (see supplementary table S1). The images were analyzed using the fMRIPrep-based Harmonized AnaLysis of Functional MRI pipeline (HALFpipe, versions 1.0.0 to 1.2.1) (53), following standardized ENIGMA protocols for functional imaging, including motion, slice-timing and susceptibility-distortion correction, spatial normalization, ICA-AROMA and aCompCor denoising, grand-mean scaling, and 6-mm FWHM smoothing. (see http://enigma.ini.usc.edu/protocols/functional-protocols/).

ROIs comprised 400 cortical regions (Schaefer-400, 17 networks), 17 subcortical (Harvard–Oxford) and 17 cerebellar (Buckner-17) regions. After excluding ROIs with <80% voxel coverage, four 6-mm spherical amygdala and accumbens ROIs (NeuroSynth peak coordinates) were added, yielding 318 regions. Time series were extracted and high-pass filtered (Gaussian-weighted, 125-s width).

### Topological data analysis and Hodge Laplacian

To capture higher-order organization that pairwise FC cannot represent, we modelled each FC matrix as a simplicial complex restricted to its 1-skeleton (nodes and edges), so that only connected components (0-cycles) and loops (1-cycles) arise. Higher-dimensional structures are likely to be unstable in brain networks (49). Loops of the network correspond to the harmonic component (kernel) of the 1-dimensional Hodge Laplacian,

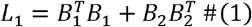

and the harmonic component is isomorphic to the first homology group. Therefore it gives an algebraic representation of the network’s 1-cycles (54). Full definitions of the boundary matrices, filtration, and Hodge decomposition are given in the Supplementary Methods.

Because a whole-brain graph contains many interlocking loops, the raw harmonic solution is spatially diffuse and cannot be localized to specific connections. To obtain a localized, interpretable basis, we combined the the Hodge Laplacian with a graph-based persistent homology filtration (see Supplementary Methods). In short, sequentially removing edges in order of FC strength separates the network into a maximum spanning tree (MST) and off-tree edges, each of which closes exactly one fundamental 1-cycle (23). This yields *Q* = m – (n – 1) linearly independent cycles for a network of *n* nodes and *m* edges, with each cycle localized to specific edges. Collecting these eigenvectors from the group-averaged FC matrix yields a common 1-cycle basis *ϕ* = [*C*_l_, *C*_2_,…, *C*_*Q*_ ].

To quantify how strongly each subject expresses the canonical cycle basis, the vectorized upper-triangular FC of each subject *w* is projected onto it,

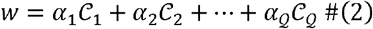

with coefficients estimated using least-squares approach as:

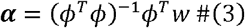

Group differences of 1-cycle coefficients are then determined using the permutation test.

### Characterization of OCD brain networks with the Hodge Laplacian

For each participant, we derived a 318 × 318 resting-state FC matrix (Pearson’s correlations with Fisher Z-transformation), vectorized the upper-triangular entries. A common basis of *Q* = 50,086 harmonic 1-cycles in an edge space of dimension *m* = 50403 was extracted from the group-averaged FC matrix across all participants, serving as a shared template, and every subject’s FC was projected onto it to yield a 2,052 × 50,086 cycle-coefficient matrix. For each α_k_ in the coefficient vector *α*, group differences were tested with a GLM including diagnosis as the effect of interest and age, sex, mean framewise displacement (FD), and scan site (fixed-effect) as covariates. We used feature-level adjustment rather than data harmonization (e.g., ComBat) because altering raw connectivity would disrupt the edge-weight rank order and distort the filtration. To incorporate the GLM model in the permutation framework, we used the Freedman-Lane permutation method (35). Briefly, residuals of a nuisance-only model were permuted, stratified by site to preserve exchangeability, and recombined to build the null. Across 50,000 permutations, FWER correction compared each cycle’s T statistic to the per-permutation maximum. Effect sizes (Cohen’s *d*) were derived from the GLM parameters, normalized by the residual SD. Significant 1-cycles (global FWER < 0.05) were assigned network-level profiles: each edge was labelled by one of 55 functional-interaction categories across 10 networks, and each cycle was binarized and summarized as a normalized 55-dimensional profile. Pairwise cosine distances and agglomerative clustering (average linkage, threshold 0.5) defined cycle clusters, for which we report loop count, mean loop size, and the fraction of constituent edges with significant edge-wise FC differences.

To further examine clinical heterogeneity, OCD samples were split by age (<18 vs. ≥18 years), current medication status, severity (Y-BOCS ≤25 vs. >25), and adult age of onset (<18 vs. ≥18 years); subgroups with <10 per group were excluded. All subgroups used the primary network template to keep a fixed topological coordinate system and were compared against HC with the main-analysis framework. Medicated and unmedicated patients were compared on age, age of onset, and severity (two-sample *t* and χ^2^ tests). To test whether the same clusters recurred, each subgroup’s significant loops were assigned to the main-analysis cluster of highest cosine similarity (labelled “Mixed” if <0.30), and per family we counted loops and computed their median absolute effect size.

### Cluster-level group effects, symptom association, and site generalization

To summarize the discriminating loops at the level of the functional clusters, and to test whether their group effects were carried by all sites or driven by a few, we computed a cluster score for each participant. For participant *i* and cluster *c*, the cluster score *s*_*l,C*_ was the mean of these sign-aligned, standardized coefficients over the loops assigned to the cluster:

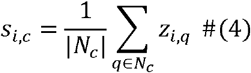

where *N*_*C*_ is the set of loops in cluster *c* and *z*_*i,q*_ is the sign-aligned, z-scored coefficient of loop q for participant *i*. Each cluster score was entered into a GLM with diagnosis as the effect of interest and age, sex, head motion, and site (fixed effect) as covariates. The standardized group effect was quantified as Cohen’s *d*. As an exploratory analysis, Y-BOCS total score was regressed on each cluster score (adjusting for age, sex, FD, site), with significance across the five clusters BH-FDR corrected. Site dependence was assessed (i) within each site (same model without the site term; sites with ≥10 participants and ≥3 per group) and (ii) under a LOSO scheme refitting the full model with each site removed. Per cluster we report the pooled effect, direction consistency across sites, the LOSO range of *d*, the most influential site, and the number of LOSO folds losing significance.

### Specificity and robustness of the 1-cycle effects

We performed several additional analyses to test the sensitivity and robustness of our method. First, we site-stratified split the 1,028 controls into two matched subgroups and repeated the full Hodge Laplacian workflow to check for spurious topology; the comparison should be null. Second, to test whether detected effects merely reflected edges with significant pairwise differences, we identified group FC differences with an LME model (group as fixed effect; age, sex, FD as covariates; site as random intercept; BH-FDR corrected), as in prior ENIGMA-OCD work on resting state functional connectome (34). Multiple comparisons correction for the FC comparison was applied using BH-FDR correction. We also analyzed the FC differences in each subgroup. qFDR for each comparison was Bonferroni corrected (0.05/9). The significant FCs were then mapped onto the cycles (green edges in figures). Third, to test whether the MST backbone drove significance, we decomposed each coefficient into tree and off-tree contributions:

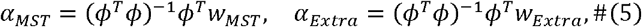

where *w*_*Extra*_ = *w* - *w*_*MST*_ . We then compared the two with paired *t*-tests over the main-analysis cycles (FDR corrected).

## Data and materials availability

The full ENIGMA OCD data are not publicly available in a repository as they contain information that could compromise the privacy of research participants. Access to these data may be requested through the ENIGMA OCD Working Group data access procedures and is subject to approval and completion of the required data-use agreements. Information about the ENIGMA OCD Working Group can be found here: https://enigma.ini.usc.edu/ongoing/enigma-ocd-working-group/. The source codes for the Hodge Laplacian analysis, sensitivity analysis, visualization, as well as non–participant-level outputs in this study are openly available at https://doi.org/10.5281/zenodo.18861978. All other data needed to evaluate the conclusions are available in the Article and/or Supplementary Information.

## Acknowledgments

The ENIGMA-Obsessive Compulsive Disorder Working-Group gratefully acknowledges support from: the Amsterdam Neuroscience (Grant No. CIA-2019-03-A to O.A.vdH., Amsterdam Neuroscience Alliance Project to G.A.vW.), the NIH Big Data to Knowledge (BD2K) award for foundational support and consortium development (Grant No. U54 EB020403 to P.M.T.). For a complete list of ENIGMA-related grant support please see here: http://enigma.ini.usc.edu/about-2/funding/. Additional funding was supported by Japan Society for the Promotion of Science (KAKENHI Grant No. 23K14825 to Y.A., No. 19K03309, 23K02956, 23K070004, 23K22361, 24K21493, 24K06547, 25K00879, 25K06842, 25H01085 to Y.H.); Carlos III Health Institute grants (Grant No. PI18/00856 and PI25/01407 to P.A., PFIS Grant No. FI17/00294 to I.M-Z, PI19/01171 to C.S-M); Fundació La Marató (Grant No. 202201 to P.A., Grant No. 091810 to L.L.); the National Institute of Mental Health (Grant No. 5R01MH116038 to A.A., Grant No. R01MH085900 and R01MH121520 to J.D.F., Grant No. K23 MH115206 to P.G., Grant No. R21MH101441 to R.M., Grant No. R21MH093889 and R01MH104648 to R.M. and H.B.S, Grant No. R01MH085900 and R01MH081864 to J.O., Grant No. K24MH121571 to C.P., Grant No. R01MH126981, R01MH111794 and R33MH107589 to E.R.S., Grant No. R01MH116147, P41EB015922, and R01MH123163 to P.M.T.); DHR-ICMR Young Medical Faculty PhD Program (Grant No. 2024-45 to S.B.); the Hartmann Müller Foundation (Grant No. 1460 to S.B.); the International Obsessive-Compulsive Disorder Foundation (IOCDF) Research Award (to P.G.); the Japan Agency for Medical Research and Development (AMED Brain/MINDS Beyond program Grant No. JP22dm0307002 to Y.H., Grant No. JP23wm0625001 and JP24wm0625204 to Y.S.); Michael Smith Foundation for Health Research (to F.J-F.); the Korea Brain Research Institute (KBRI, the Brain Science Convergence Research Program RS-2023-00266120 and Basic Research Program 25-BR-05-05 to M.K.); the Spanish Ministry of Health (Fondo de Investigaciones Sanitarias, project P11/01419 to L.L.); the Medical Research Council of South Africa (SAMRC; to C.L.); the National Research Foundation of South Africa (to C.L.); the Portuguese Foundation for Science and Technology (Fundação para a Ciência e a Tecnologia; Grant No. UIDB/50026/2020 and UIDP/50026/2020 to P.M.); the Norte Portugal Regional Operational Programme (NORTE 2020; Grant No. NORTE-01-0145-FEDER-000013 and NORTE-01-0145-FEDER-000023 to P.M., under the PORTUGAL 2020 Partnership Agreement, through the European Regional Development Fund); the FLAD Science Award Mental Health 2021 (to P.M.); the Government of India grants from the Department of Science and Technology (DST INSPIRE faculty Grant No. IFA12-LSBM-26 to J.C.N., Grants No. SR/S0/HS/0016/2011 to J.Y.C.R.) and from the Department of Biotechnology (Grant No. BT/06/IYBA/2012 to J.C.N., Grant No. BT/ PR13334/Med/30/259/2009 to J.Y.C.R.); the Ministry of Science, Innovation and Universities and the European Union—NextGenerationEU (Grant No. RYC2021-031228-I to M.P-P.); Italian Ministry of Health (Ricerca Corrente 2022, 2023 to F.P.); Department of Biotechnology - Wellcome Trust India Alliance (Early Career Fellowship Grant No. IA/CPHE/18/1/503956 to V.S., Grant No. IA/CRC/19/1/610005 and Senior Fellowship Grant No. 500236/Z/11/Z to G.V.); the Catalan Agency for the Management of University and Research Grants (AUGUR Grant No. 2017SGR 1247 to C.S-M.); the Helse Vest Health Authority (Grant No. 911754 and 911880 to A.L.T.); the National Institute on Aging Research Project Grant Program (Grant No. R01AG058854 to Y.D.vdW.); the ENIGMA Parkinson’s Initiative: A Global Initiative for Parkinson’s Disease, NINDS award (RO1NS107513 to Y.D.vdW.); Dutch Organization for Scientific Research (NWO/ZonMW) VENI grant (No. 165.610.002 and 016.156.318 to W.vL. and G.vW., No. 917.15.318 to G.vW., No. 916-86-038 to O.A.vdH.); the Swiss National Science Foundation (Grant No. 320030_130237 to S.W.); the National Natural Science Foundation of China (Grant No. 82071518 to Z.W.); the Obsessive-Compulsive Foundation (to D.J.S.); the Brain & Behavior Research Foundation (NARSAD grant) and Netherlands Brain Foundation (Grant No. 2010(1)−50 to O.A.vdH.); the Deutsche Forschungsgemeinschaft (DFG; Grant No. KO 3744/11-1 to K.K.)

## Author contributions

H.R., M.K.C, O.A.vdH., D.J.S., P.M.T., K.K. designed research; H.R. performed research; M.K.C., D.J.S., O.A.vdH., P.M.T., K.K. supervised the research; H.R., M.K.C., W.B.B., N.D., Y.A., P.A., A.A., S.B., M.C.B., F.B., S.B., S.B., Y.C., F.C., B.C., G.K.E., S.F., J.D.F., R.G.G., P.G., K.H., B.H., Y.H., M.Q.H., J.I., F.J-F., M.K., J.S.K., L.L., C.R.L., C.L., R.M., I.M-Z., J.M.M., P.S.M., P.M., E.M., A.N., J.C.N., E.L.N., J.O., J.C.P., J.C.P., M.P-P., F.P., F.P., C.P., J.Y.R.C., D.R.M., Y.S., J.R.S., E.S., V.S., B.H.S., C.S-M., N.S., E.R.S., E.S., P.R.S., S.I.T., A.L.T., B.V., A.vsS., Y.D.vdW., W.vL., H.vM., G.vW., D.V., G.V., C.V., S.W., Z.W., T.Y., J.Y.Y., Q.Z., P.M.T., D.J.S., O.A.vdH., K.K. collected data for this study; H.R. wrote the paper.

