## Supplementary Materials for "Mapping Higher-Order Topology in OCD Brain Networks with Hodge Laplacian"

**Supplementary Methods**

**Subject exclusion**

The exclusion process was kept the same with the previous ENIGMA-OCD connectome research. Specifically, we excluded 2 healthy controls who were using psychotropic medication, 264 participants whose data failed neuroimaging quality control, 111 participants due to excessive motion, 315 participants with insufficient brain coverage and 151 participants from samples with <10 participants per group. Finally, 1024 OCD patients and 1028 healthy controls, in total 2052 subjects were included in the study.

**Spectral graph theory**

A FC matrix of brain network can be represented as a weighted undirected complete graph $G=(V, E, w)$, where the vertex set $V$ corresponds to ROIs, edge set$E$ encodes FCs and w represents edge weights. In spectral graph theory, the graph $G$ is represented by its adjacency matrix and Laplacian matrix. The adjacency matrix $A$ contains the functional coupling between ROIs, and can be expressed as:

$\begin{aligned} A\left( i,j \right)= \left\{ \begin{aligned} w_{ij}, \left( v_{i}, v_{j} \right)\in E \\ 0,{(v}_{i}, v_{j})\notin E \end{aligned} \right. \#\left( 1 \right) \end{aligned}$

The Laplacian matrix is defined as $L_{0}=D-A$, where $D$ is the degree matrix of $G$ with its diagonal elements being the degrees of nodes (1). Specifically, the $L_{0}$ matrix can be expressed as:

$$\begin{aligned} L_{0}\left( i,j \right)= \left\{ \begin{aligned} \deg\left( v_{i} \right), i=j \\ -1, i\neq j and \left( v_{i}, v_{j} \right)\in E \\ 0,{i\neq j and (v}_{i}, v_{j})\notin E \end{aligned} \right. \#\left( 2 \right) \end{aligned}$$

$L_{0}$ is considered as a central operator in spectral graph theory. Its spectrum captures the fundamental features of the network’s structure, including connectedness and the presence of modular organization. Importantly, the number (multiplicity) of zero eigenvalues of $L_{0}$ corresponds to the number of connected components of the graph $G$. The proposed Hodge Laplacian generalizes $L_{0}$ from graphs to simplicial complexes, enabling spectral analysis of higher-dimensional topological structures such as cycles.

**Simplicial complex and chain complex**

While the graph representation $G=(V, E, w)$ captures pairwise functional couplings, many organizational features of brain networks arise from higher-order multi-nodal interactions that cannot be described solely by edges. This can be achieved by generalizing the graph into the simplicial complex as its higher-order representation. For the $k$-th dimension, a $k$-simplex $\sigma^{k}=\left\{ v_{0},v_{1}, \ldots, v_{k} \right\}$ is a finite set of $k+1$ distinct vertices. Geometrically, for instance, a 0-simplex is a vertex which will be a point/node, 1-simplex is an edge with 2 vertices, and a 2-simplex is a triangle with 3 vertices. A simplicial complex $K$ is then a collection of simplices up to a certain dimension.

In order to calculate the simplicial complex, one needs to define the orientation of a simplex that belongs to any above-0 dimension. Defining an orientation is necessary as signed edges are required for linear algebra operations to be applied. The orientation is usually specified according to the ordering of the vertices, either ascending or descending. For example, a 1-simplex $\sigma^{1}=\left\{ v_{0},v_{1} \right\}$ is an edge and can be denoted as either $v_{0}- v_{1}$ or $v_{1}- v_{0}$, which means its direction is from $v_{1}$ to $v_{0}$ or from $v_{0}$ to $v_{1}$. With oriented $k$-simplices, we can sum them up and form a $k$-th chain group $C_{k}$. In other words, once every simplex has a direction (orientation), we can treat them as if they are signed building blocks and take linear combinations of them. All such combinations together form a vector space called the $k$-th chain group, denoted $C_{k}$. It is important to note that these assigned directions only represent higher-order statistical dependencies, rather than direct, causal neuronal transmission in the context of functional brain networks.

A chain complex is then defined as a sequence of these chain groups and the boundary operators are used for connecting chain groups from different dimensions (2). For an oriented $k$-simplex, the boundary operator $\partial_{k} : C_{k}\to C_{k-1}$ is defined as:

$$\begin{aligned} \partial_{k}\sigma^{k}=\sum_{i=0}^{k} \left( -1 \right)^{i}\sigma_{i}^{k-1}= \sum_{i=0}^{k} \left( -1 \right)^{i}\left[ v_{0}, v_{1}, \ldots,\hat{v_{i}}, \ldots,v_{k} \right] \#\left( 3 \right) \end{aligned}$$

where $\left[ v_{0}, v_{1}, \ldots,\hat{v_{i}}, \ldots,v_{k} \right]$ is an oriented $(k-1)$-simplex generated from the vertices inside $\sigma^{k}$ except $\hat{v_{i}}$. A simplex can be mapped to its boundaries by the boundary operator, for instance, an edge can be mapped to its two vertices, and a triangle to its three edges. This operation also guarantees that $\partial_{k-1}\partial_{k}=0$, indicating that the boundary of a boundary is empty. Thus, a chain complex can be expressed as:

$$\begin{aligned} \ldots\underset{\to}{\partial_{k+1}}C_{k}\left( K \right)\underset{\to}{\partial_{k}}C_{k-1}\left( K \right)\underset{\to}{\partial_{k-1}}\ldots\underset{\to}{\partial_{1}}C_{0}\left( K \right)\underset{\to}{\partial_{0}}0\#\left( 4 \right) \end{aligned}$$

Here, the boundary of the 0th chain group is an empty set. Also, for any n that exceeds the maximum dimension $k$, $C_{n}\left( K \right)$ is an empty vector space and the corresponding boundary operator is a zero map. See Supplementary Figure S1 for an illustration. Overall, in this way, we can organize simplices by dimension into vector spaces, and connect those spaces with boundary maps that encode how higher-dimensional structures attach to lower-dimensional ones.

From a geometric perspective, the topological feature of dimension $k$ (which is usually referred to as $k$-dimensional hole) describes a closed structure that encloses a “void” and cannot be "filled in" or continuously contracted to a point within the space. For instance, a 1-dimensional hole manifests as a closed loop that is connected but surrounds an empty interior; it remains a distinct feature because it cannot be collapsed without altering the space's topology. Algebraically, these features are formalized as homology groups, defined by the relationship between $k$-cycles and $k$-boundaries. Mathematically, this geometric intuition is formalized as an algebraic construction: a structure that is closed but does not bound a higher-dimensional volume. For a chain complex $C_{k}$, we distinguish two fundamental subspaces. The collection of $k$-cycles $Z_{k}$ is the kernel subspace, which is expressed as:

$$\begin{aligned} Z_{k}=\ker\partial_{k}= \left\{ \sigma^{k}\in C_{k}|\partial_{k}\sigma^{k}=0 \right\} \#\left( 5 \right) \end{aligned}$$

$Z_{k}$ contains all possible candidates of $k$-cycles, as long as they are a closed (the boundary is zero). The collection of $k$-boundaries $B_{k}$ is the image subgroup, which is expressed as:

$$\begin{aligned} B_{k}=img \partial_{k+1}= \left\{ \sigma^{k}\in C_{k}|\sigma^{k}=\partial_{k+1}\sigma^{k+1},\sigma^{k+1} \in C_{k+1} \right\} \#\left( 6 \right) \end{aligned}$$

This “closed-but-not-filled” condition implies that the cycle cannot be trivialized by the boundary operator. Specifically, a topological feature corresponds to a $k$-cycle (an element of $\ker\partial_{k}$) that fails to be a $k$-boundary (an element of $img \partial_{k+1}$). Consequently, the $k$-homology class $H_{k}$ is then represented as $H_{k}= Z_{k}/B_{k}$. Importantly, $H_{k}$ is not a single cycle but an equivalence class of cycles. As long as the difference between two cycles is a boundary, they are considered as equivalent. In this study, as we only focus on the graph with the highest dimension of 1, a 0-dimensional cycle is then a connected component or a node, and a 1-dimensional cycle is a loop.

**Persistent homology and graph filtration**

PH provides a framework for tracking how topological features—such as connected components, loops—emerge and disappear as a resolution or distance threshold varies. In classical PH, a sequence of nested simplicial complexes, which usually refers to a filtration, is constructed (3). The birth and death of $k$-dimensional features along this sequence are quantified using the duration of filtration values from birth to death known as persistence (4). The persistence is usually represented as one-dimensional intervals in a persistent barcode. Although the present study did not compute full persistence, the underlying concept of a filtration is essential for finding all possible 1-cycles in the graph.

For a weighted undirected complete graph $G$ with $m$ number of edges, the filtration process starts with the complete graph and is a sequence of sub-graphs ${{(G}_{\epsilon_{t}})}_{t=1}^{m}$ of $G$:

$$\begin{aligned} G\subset G_{\epsilon_{m}}\subset\ldots\subset G_{\epsilon_{2}}\subset G_{\epsilon_{1}} \#\left( 7 \right) \end{aligned}$$

where $\epsilon_{1}<\epsilon_{2}<\ldots<\epsilon_{m}$ are the sorted edge weights from the graph (5, 6). Every subgraph is generated by removing one edge with the highest edge weights from the previous subgraph.

**Birth-death decomposition**

Here we present a toy model of a 4-node graph as an example (Supplementary Figure S2). In the context of persistent homology, the evolution of topological features is typically quantified by Betti numbers, where $\beta_{0}$ represents the number of connected components and $\beta_{1}$ represents the number of cycles (loops), as demonstrated in the visualization. The red barcodes represent the 0-cycles or the connected components in the graph. Notice that once a component is born, it does not die, so all the connected components have $\infty$ death values which can be ignored. For a graph with $n$ nodes, the total number $\mathcal{P}$ of birth values of connected components is $n-1$, which correspond to the $\beta_{0}$ number of the graph. The light red barcode which corresponds to the original complete graph is taken out because it doesn’t have a birth value. The birth value set $\mathcal{B}\left( G \right)$ of the graph $G$ is then an increasing set of edge weights. The green barcodes represent the 1-cycles or the loops in the graph. All the loops are naturally existing in the complete graph; thus, all the birth values are $-\infty$ and can be ignored. The death value set $\mathcal{D}\left( G \right)$ of the graph $G$ is then a decreasing set of edge weights. The two sets together compose all edges in the graph since they cover all filtration values in the filtration process, thus we have:

$$\begin{aligned} m=\frac{n\left( n-1 \right)}{2}\mathcal{=P+ Q \#}\left( 8 \right) \end{aligned}$$

The total number $\mathcal{Q}$ of death values of loops is then $(n-1)(n-2)/2$, which correspond to the $\beta_{1}$ number of the graph.

Importantly, adding any off-tree edge back to the MST creates a unique fundamental 1-cycle. This decomposition provides the basis for constructing the 1-cycle basis. With the graph filtration, we are able to analyze every possible 1-cycle structure, and also avoid an arbitrary connectivity threshold which is one of the major drawbacks of conventional graph analysis in neuroimaging (7, 8).

**Spectral simplicial complex and Hodge Laplacian**

The graph Laplacian $L_{0}$, which applied on the classical graph theory analysis with nodal interactions, can be viewed as the 0-dimensional instance of a more generalized framework that extends spectral graph theory to simplices or higher-order interactions. This generalization is provided by the Hodge Laplacian, which is defined on the chain complex (9, 10). While the boundary operator $\partial_{k}$ defines the topological relationships algebraically, its implementation for data analysis requires a matrix form. For a chain complex $C$, its $k$-th boundary matrix $B_{k}$ is defined as:

$$\begin{aligned} B_{k}\left( i,j \right)= \left\{ \begin{aligned} 1, if \sigma_{i}^{k-1}\subset\sigma_{j}^{k} and \sigma_{i}^{k-1}\sim\sigma_{j}^{k} \\ -1, if \sigma_{i}^{k-1}\subset\sigma_{j}^{k} and \sigma_{i}^{k-1}≁\sigma_{j}^{k} \\ 0, if \sigma_{i}^{k-1}\not\subset\sigma_{j}^{k} \end{aligned} \right. \#\left( 9 \right) \end{aligned}$$

Same as the boundary operator, the boundary matrices also satisfy that $B_{k}B_{k+1}=0$. The $k$-th Laplacian matrix can be expressed as:

$$\begin{aligned} {L_{k}=B_{k}^{T}B}_{k}+B_{k+1}B_{k+1}^{T} \#\left( 10 \right) \end{aligned}$$

There is no boundary for 0-dimensional simplices ($B_{0}=0$), the 0-th Laplacian, which is the graph Laplacian $L_{0}= B_{1}B_{1}^{T}$. As the highest dimension of the graph is 1, the 2-nd boundary matrix on a graph-based simplicial complex is also zero. Thus the 1-st Laplacian is ${L_{1}=B_{1}^{T}B}_{1}$.

**Hodge decomposition and algebraic representation of 1-cycles**

In order to identify and perform statistical comparisons for $k$-cycles, we first need to perform the Hodge decomposition. A $k$-th chain group will be then decomposed into three orthogonal subspaces:

$$\begin{aligned} C_{k}\left( K \right)=imgB_{k+1}\oplus\ker L_{k}\oplus imgB_{k}^{T} \#\left( 11 \right) \end{aligned}$$

The gradient component corresponds to chains induced by lower-dimensional potentials, the curl component consists of boundaries of higher-dimensional simplices, and the harmonic component comprises divergence-free and curl-free chains. The harmonic subspace provides a direct mathematical representation of non-trivial topological cycles, which are isomorphic to the $k$-th homology group (11). In this study, we mainly focus on the second subspace, which is $ker L_{k}$. Statistical analyses were performed on this harmonic component to isolate genuine $k$-cycle differences across subjects. First, we solve the eigen-decomposition of $L_{k}$:

$$\begin{aligned} L_{k}=U_{k}\Lambda_{k}U_{k}^{T} \#\left( 12 \right) \end{aligned}$$

where $U_{k}$ is the matrix of eigenvectors, and k is a diagonal matrix of eigenvalues. Here, as the generalization of spectral graph theory, the multiplicity of zero eigenvalues of $L_{k}$ is equal to the $k$-th Betti number $\beta_{k}$. Harmonic eigenvectors corresponding to the zero eigenvalues can be expressed in the basis of $k$-simplices. The numerical entries of the eigenvector are therefore the coefficients of these simplices in the corresponding simplest possible $k$-cycle. A non-zero coefficient indicates that the simplex contributes to the cycle, whereas a zero coefficient indicates non-participation. Thus, the pattern of non-zero entries in a harmonic eigenvector can help to localize which simplices form the underlying topological feature. Given a $k$-cycle $\mathcal{C}_{k}$ that belongs to the $k$-th chain group $C_{k}$, its algebraic representation can be expressed as:

$$\begin{aligned} \mathcal{C}_{k}=\sum_{i=1}^{j} a_{i}\sigma_{i}^{k} \#\left( 13 \right) \end{aligned}$$

where $a_{i}\mathbb{\in R}$ is the $i$-th simplex coefficient, $\sigma_{i}^{k}$ is the $i$-th oriented $k$-simplex, and the eigenvector of $\mathcal{C}_{k}$ has $j$ number of entries.

Figures

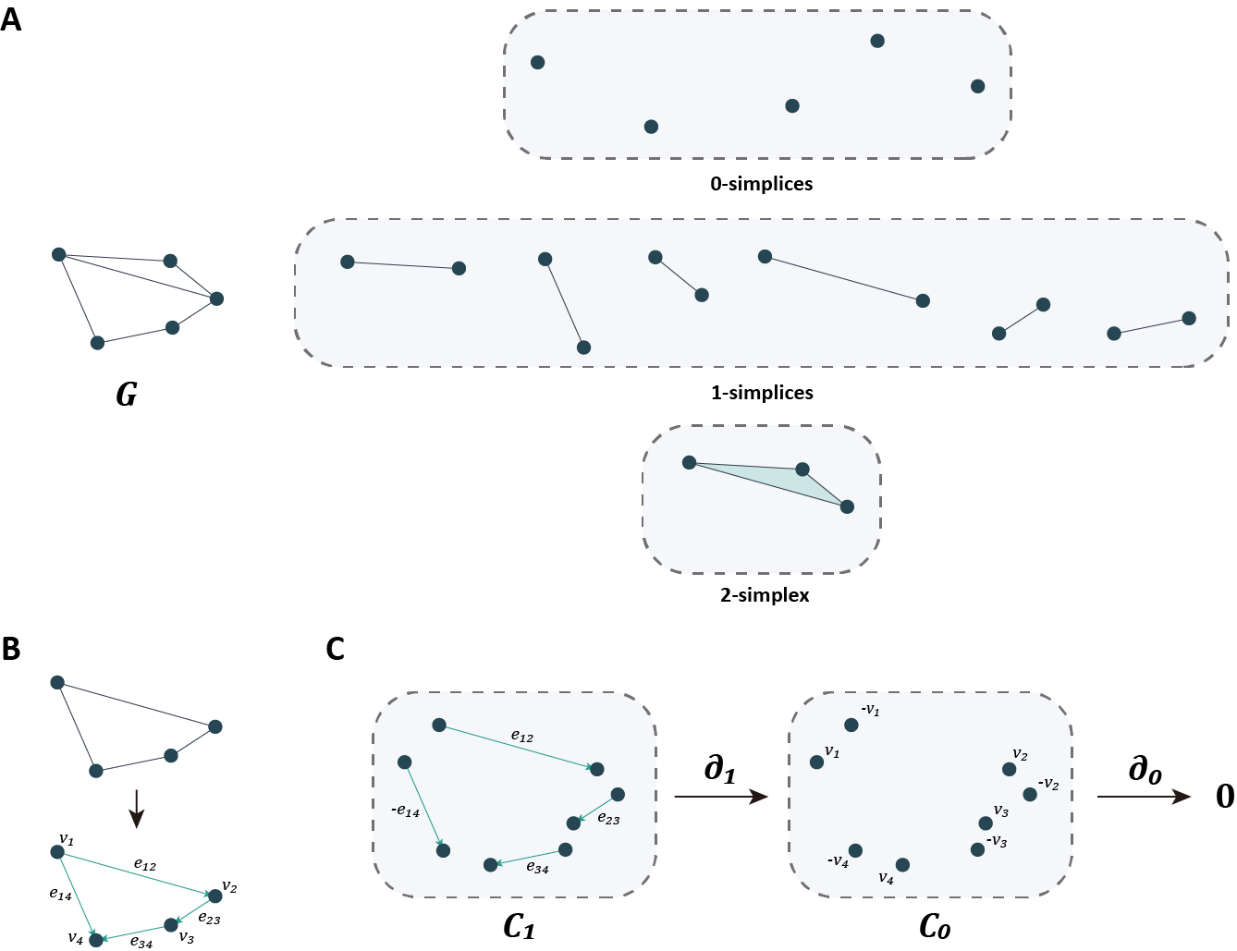

**Fig. S1. Explanation of simplicial complex, chain complex and boundary operator. A.** A toy graph $\boldsymbol{G}$ and its associated simplices: 5 vertices (0-simplices), 6 edges (1-simplices), and 1 filled triangle representing a 2-simplex. **B.** For computation and the construction of chain complex, orientation is assigned to simplices to generate oriented chains. **C.** The resulting chain groups $\boldsymbol{C}_{\boldsymbol{1}}$ and $\boldsymbol{C}_{\boldsymbol{0}}$ and the action of the boundary operators $\boldsymbol{\partial}_{\boldsymbol{1}}$ (mapping oriented edges to signed endpoints) and $\boldsymbol{\partial}_{\boldsymbol{0}}$ (mapping 0-chains to the zero element).

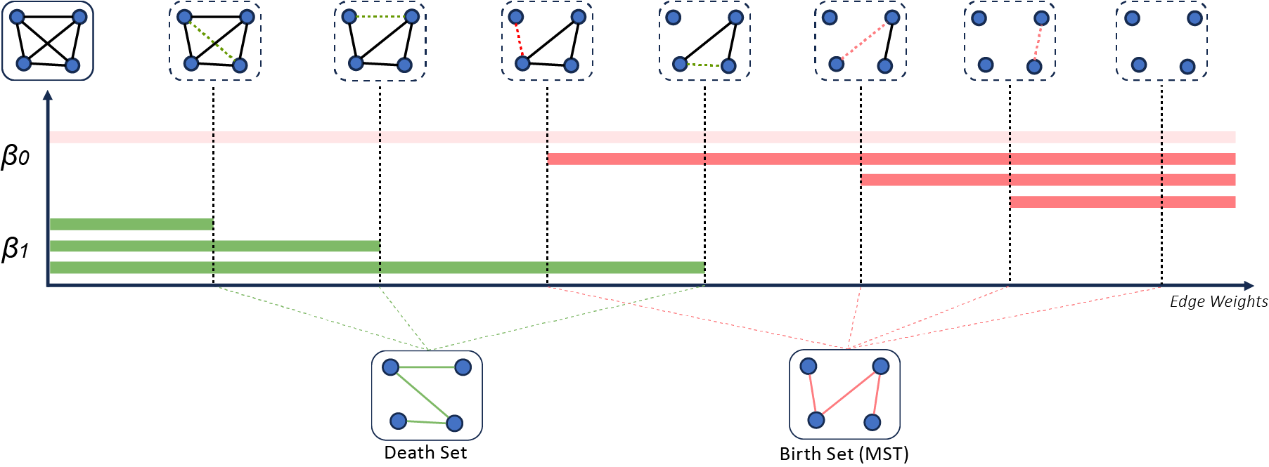

**Fig. S2. Schematic of graph filtration process.** Based on the concept of persistent homology, we perform the graph filtration according to sorted edge weights. The number of connected components ($\boldsymbol{\beta}_{\boldsymbol{0}}$) and the number of cycles ($\boldsymbol{\beta}_{\boldsymbol{1}}$) change at different filtration values. Notice that since a component has infinite death value and a cycle has infinite birth value, we can decompose edges into death set and birth set, and the birth set is exactly the maximum spanning tree (MST) structure of the graph.

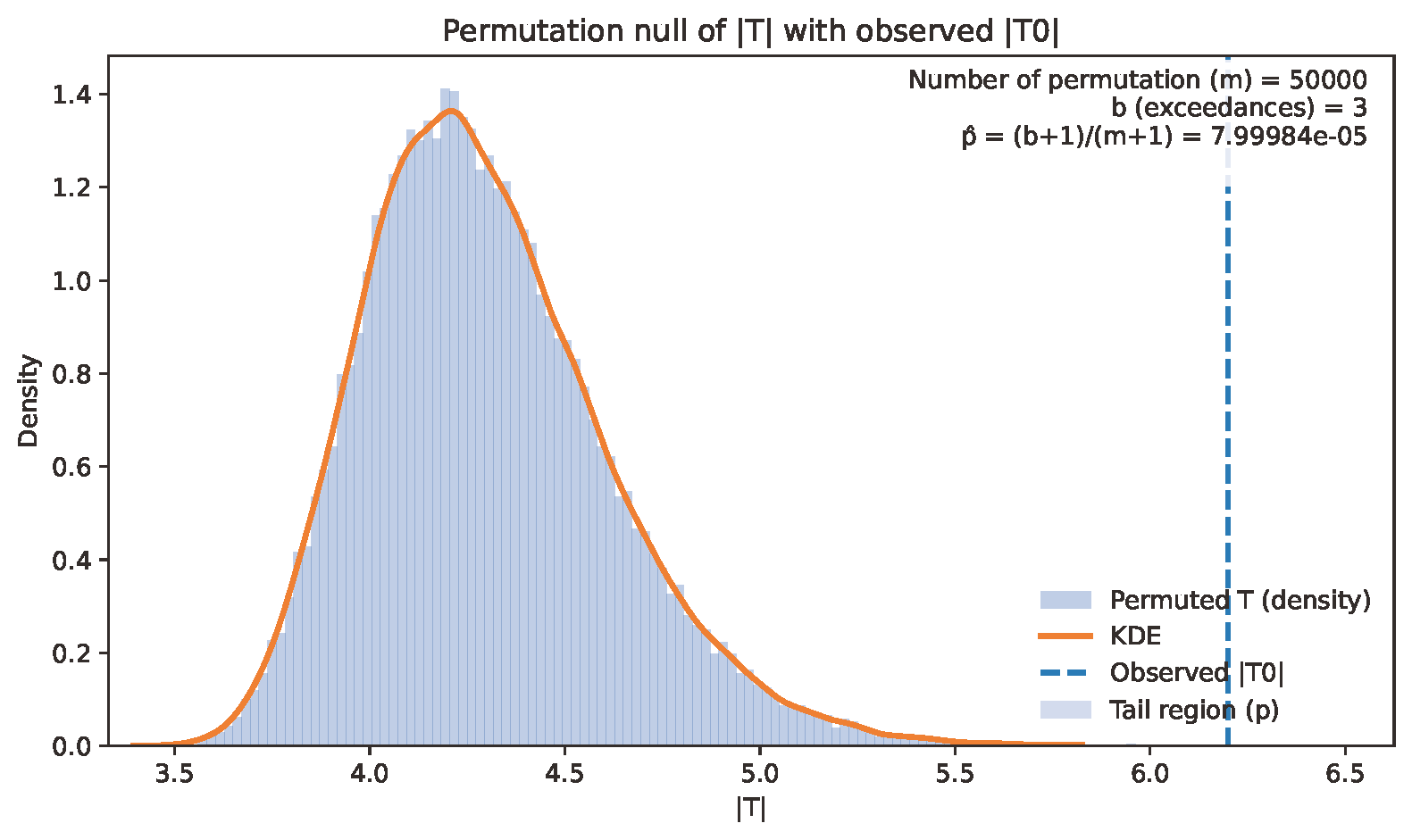

**Fig. S3.** **Null distribution of the permutation test.** The histogram and the kernel density estimation (KDE, orange curve) represent the empirical null distribution of the maximum absolute T-statistics obtained from 50,000 permutations from the main analysis. The vertical dashed line indicates the observed maximum absolute T-statistic ($\mathbf{|T0|}$). Only 3 out of 50,000 permutations exceeded the observed value, yielding a highly conservative $\boldsymbol{p}$-value of $\boldsymbol{p}\boldsymbol{\approx8\times}\mathbf{10}^{\mathbf{-5}}$. This demonstrates that the significance of discriminating 1-cycles were highly unlikely to have arisen by chance, even after controlling for multiple comparisons.

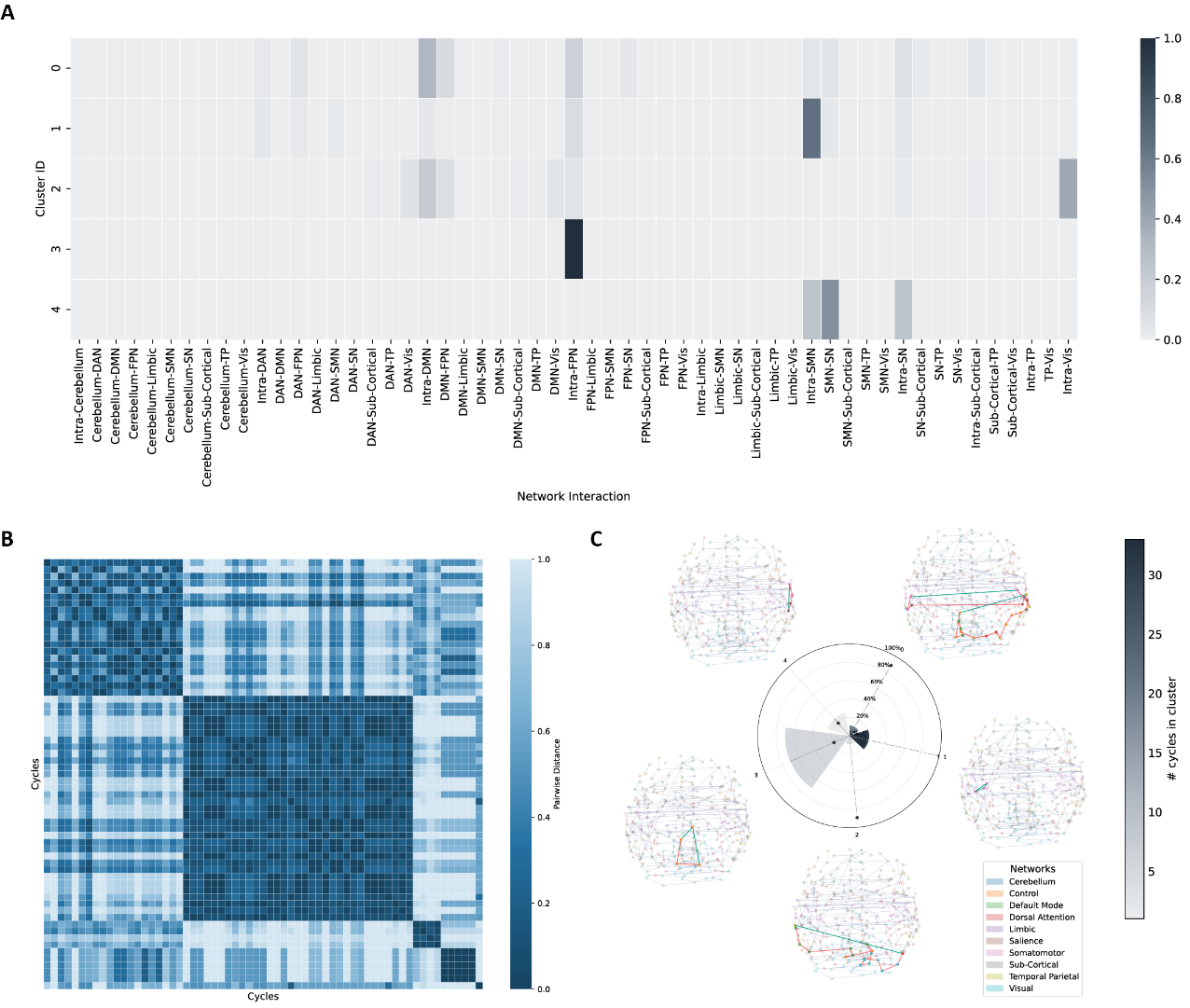

**Fig. S4.** **1-cycle abnormalities in adult sample. A.** Agglomerative clustering results for adult samples according to functional profiles of 1-cycles. **B.** Pairwise cosine distances between cycles. 5 distinct clusters are shown in the heatmap. **C.** Radial bar plot summarizing the following three cluster-level properties: Wedge height indicates the mean fraction of edges per 1-cycle that show significant edge-wise FC differences between OCD and controls, averaged across 1-cycles within each cluster. Bar color illustrates the number of significant 1-cycles in the cluster. The dot marks the cluster’s normalized mean 1-cycle length. Visualizations of the most discriminating 1-cycles in each cluster are shown around the plot. DAN: dorsal attention network, DMN: default mode network, FPN: frontoparietal network (labeled “Control” in the atlas), OCD: obsessive-compulsive disorder, SMN: somatomotor network, SN: salience network, TP: temporal parietal network, Vis: visual network.

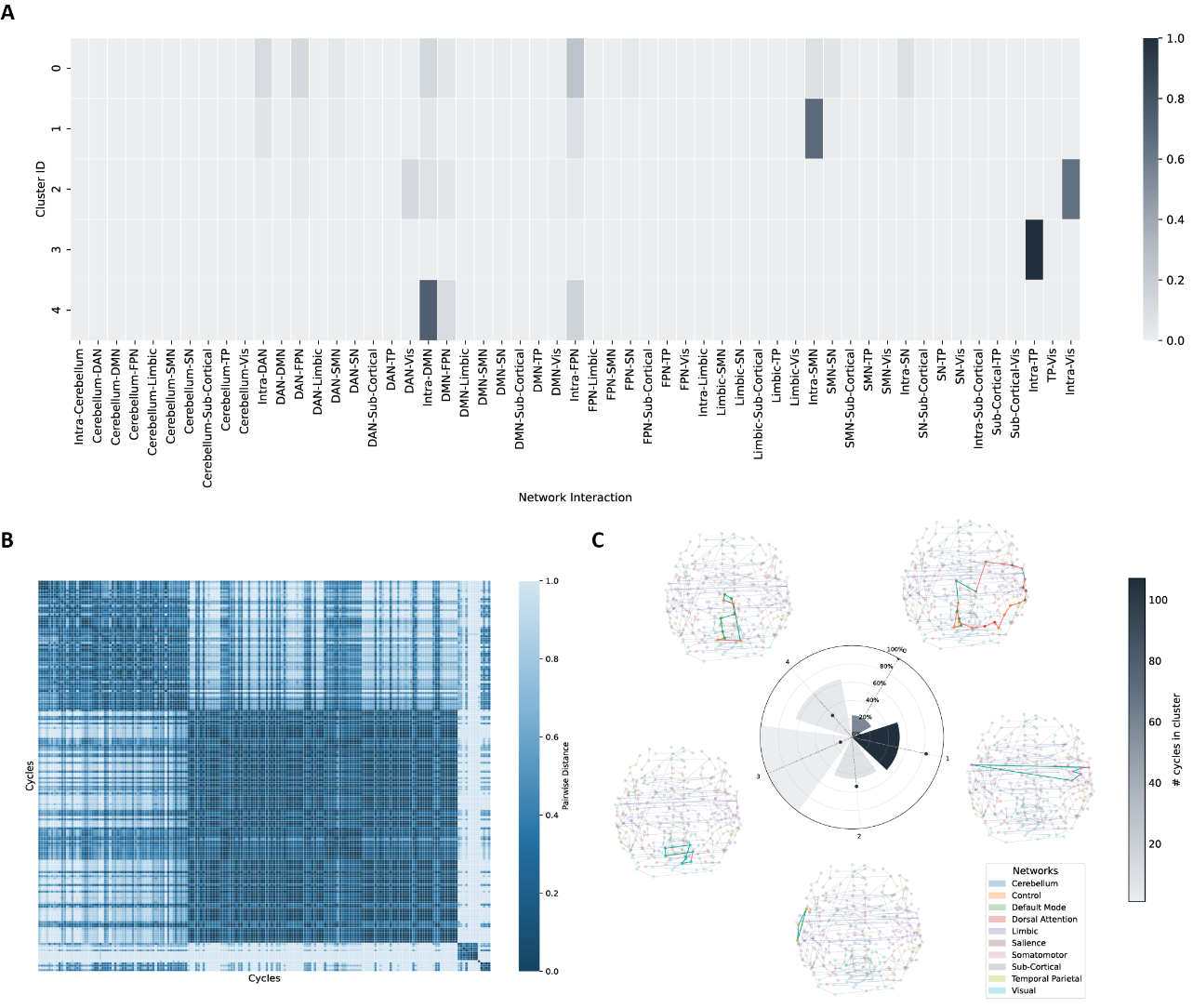

**Fig. S5.** **1-cycle abnormalities in medicated sample. A.** Agglomerative clustering results for medicated samples according to functional profiles of 1-cycles. **B.** Pairwise cosine distances between cycles. 5 distinct clusters are shown in the heatmap. **C.** Radial bar plot summarizing the following three cluster-level properties: Wedge height indicates the mean fraction of edges per 1-cycle that show significant edge-wise FC differences between OCD and controls, averaged across 1-cycles within each cluster. Bar color illustrates the number of significant 1-cycles in the cluster. The dot marks the cluster’s normalized mean 1-cycle length. Visualizations of the most discriminating 1-cycles in each cluster are shown around the plot. DAN: dorsal attention network, DMN: default mode network, FPN: frontoparietal network (labeled “Control” in the atlas), OCD: obsessive-compulsive disorder, SMN: somatomotor network, SN: salience network, TP: temporal parietal network, Vis: visual network.

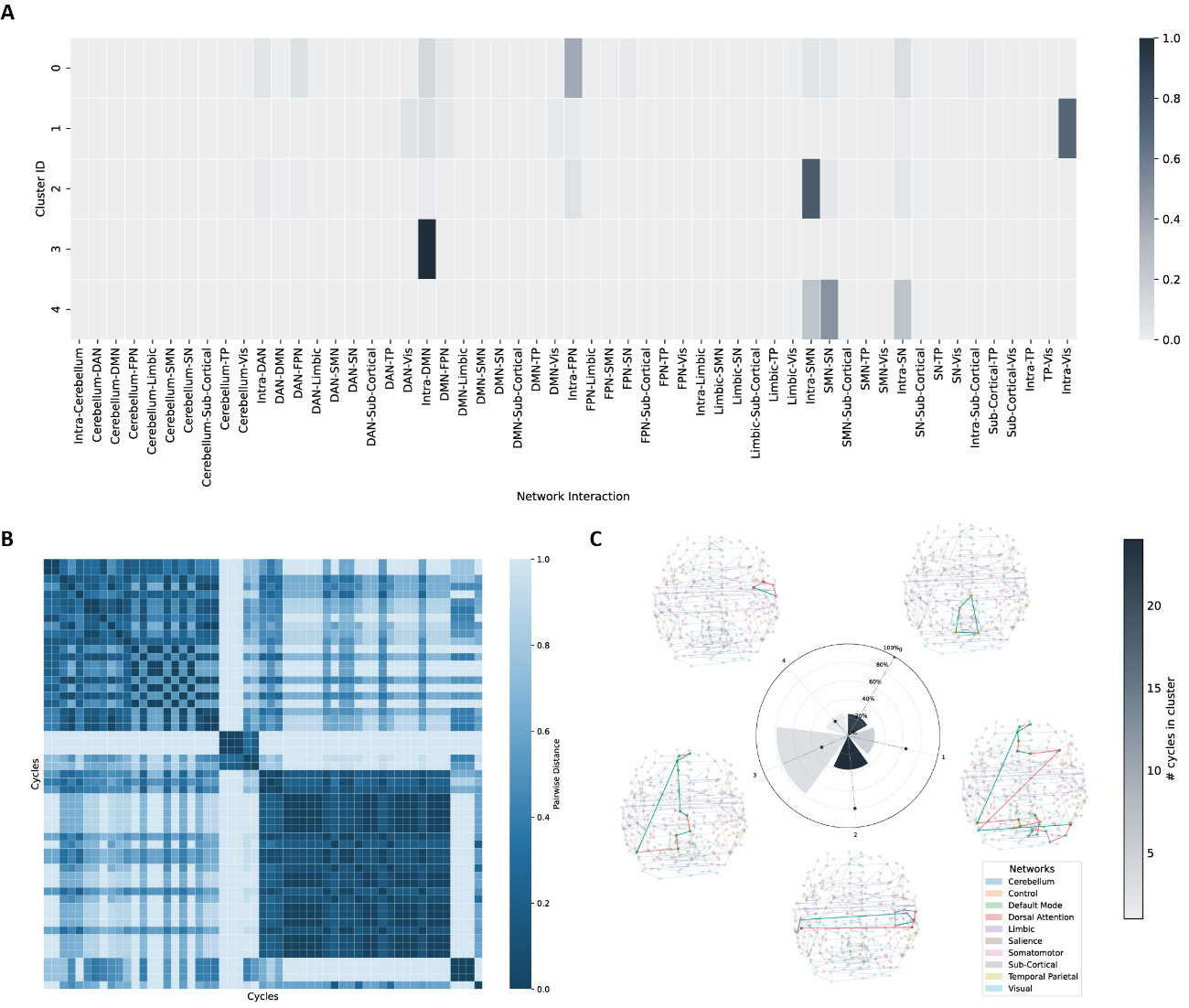

**Fig. S6.** **1-cycle abnormalities in high-severity sample. A.** Agglomerative clustering results for high-severity samples according to functional profiles of 1-cycles. **B.** Pairwise cosine distances between cycles. 5 distinct clusters are shown in the heatmap. **C.** Radial bar plot summarizing the following three cluster-level properties: Wedge height indicates the mean fraction of edges per 1-cycle that show significant edge-wise FC differences between OCD and controls, averaged across 1-cycles within each cluster. Bar color illustrates the number of significant 1-cycles in the cluster. The dot marks the cluster’s normalized mean 1-cycle length. Visualizations of the most discriminating 1-cycles in each cluster are shown around the plot. DAN: dorsal attention network, DMN: default mode network, FPN: frontoparietal network (labeled “Control” in the atlas), OCD: obsessive-compulsive disorder, SMN: somatomotor network, SN: salience network, TP: temporal parietal network, Vis: visual network.

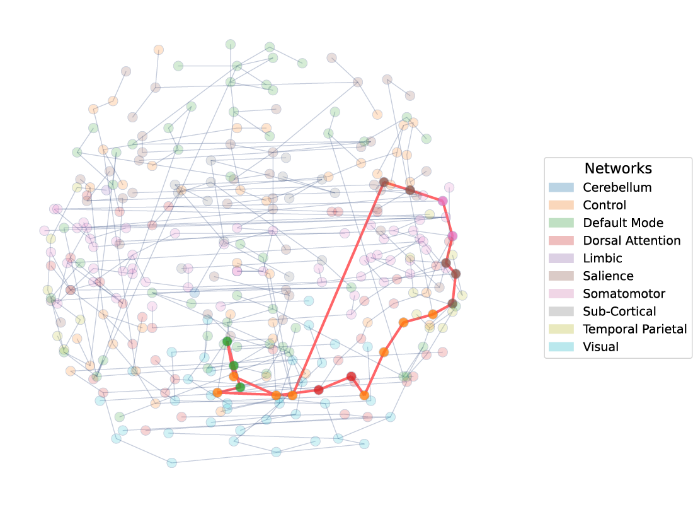

**Fig. S7.** **Discriminating 1-cycle structure of unmedicated subgroup analysis.** This 1-cycle showed interaction among default mode network, somatomotor network, dorsal attention network, salience network and control/frontoparietal network.

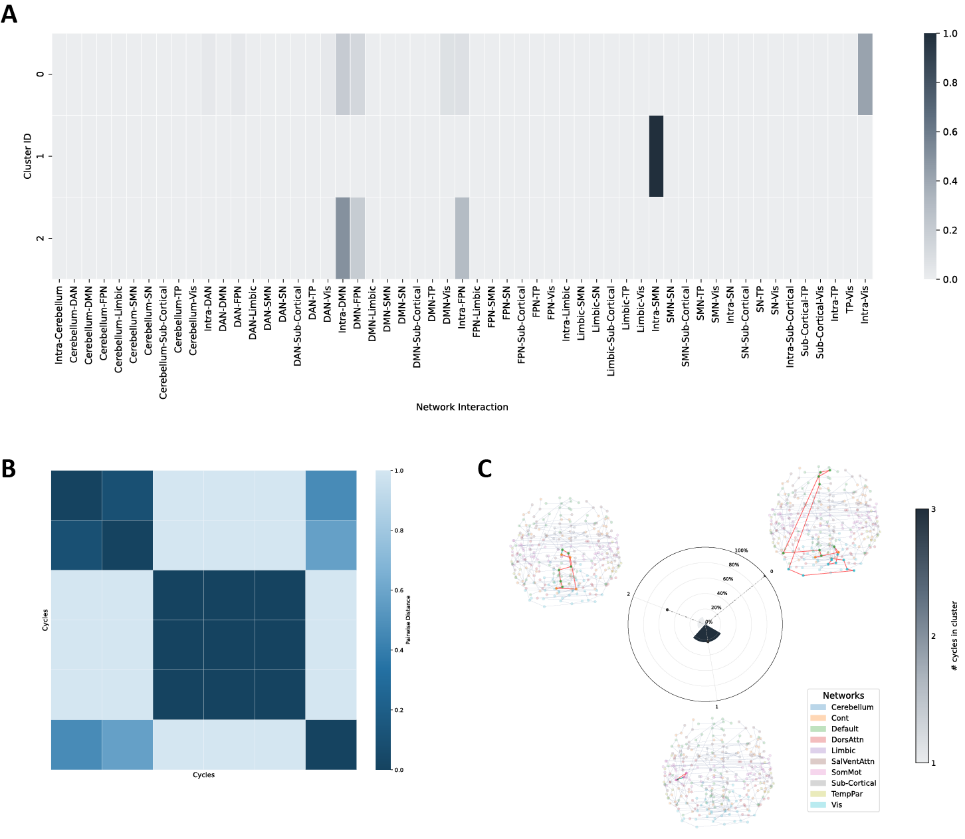

**Fig. S8.** **1-cycle abnormalities in low-severity sample. A.** Agglomerative clustering results for low severity sample according to functional profiles of cycles. 3 clusters were found in this subgroup analysis, and half of them were related to intra-SMN connections. **B.** Pairwise cosine distances between cycles. **C.** Cluster-level results summarized by a radial bar chart, where each bar represents a cluster and its height indicates the proportion of significant edges. Bar color reflects the number of 1-cycles in the cluster, and a dot marker denotes the normalized mean 1-cycle size. Visualizations of the most discriminating 1-cycles in each cluster are shown around the plot. DAN: dorsal attention network, DMN: default mode network, FPN: frontoparietal network (labeled “Control” in the atlas), OCD: obsessive-compulsive disorder, SMN: somatomotor network, SN: salience network, TP: temporal parietal network, Vis: visual network.

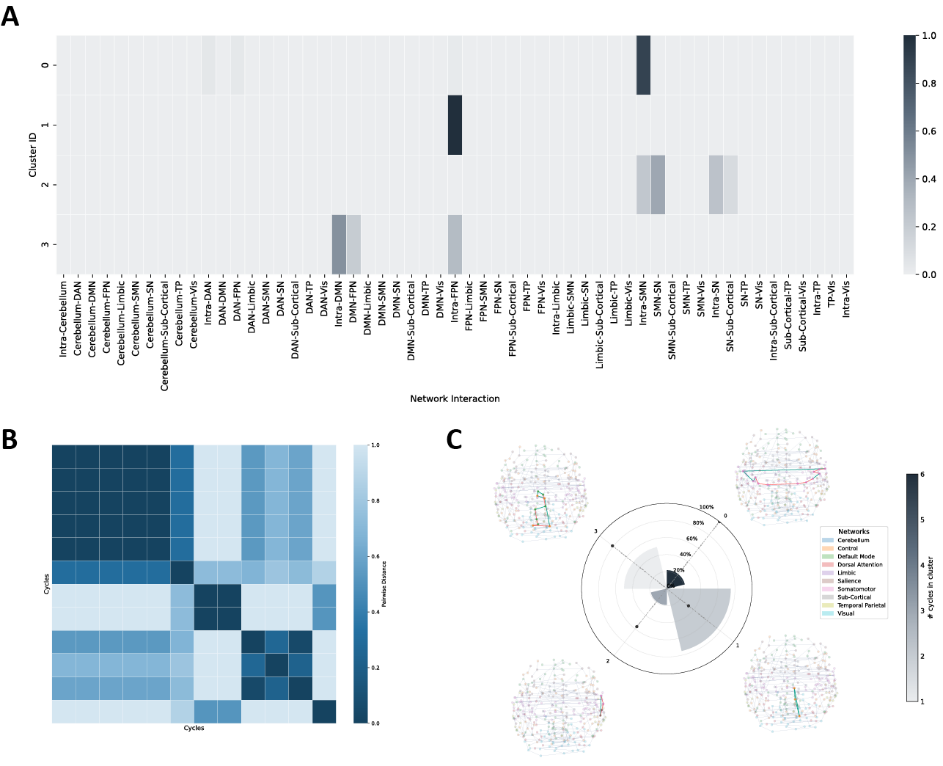

**Fig. S9.** **1-cycle abnormalities in adult-onset sample. A.** Agglomerative clustering results for adult-onset sample according to functional profiles of cycles. 4 clusters were found in this subgroup analysis, and half of them were also related to intra-SMN connections. **B.** Pairwise cosine distances between cycles. **C.** Cluster-level results summarized by a radial bar chart, and visualizations of the most discriminating 1-cycles in each cluster are shown around the plot. DAN: dorsal attention network, DMN: default mode network, FPN: frontoparietal network (labeled “Control” in the atlas), OCD: obsessive-compulsive disorder, SMN: somatomotor network, SN: salience network, TP: temporal parietal network, Vis: visual network.

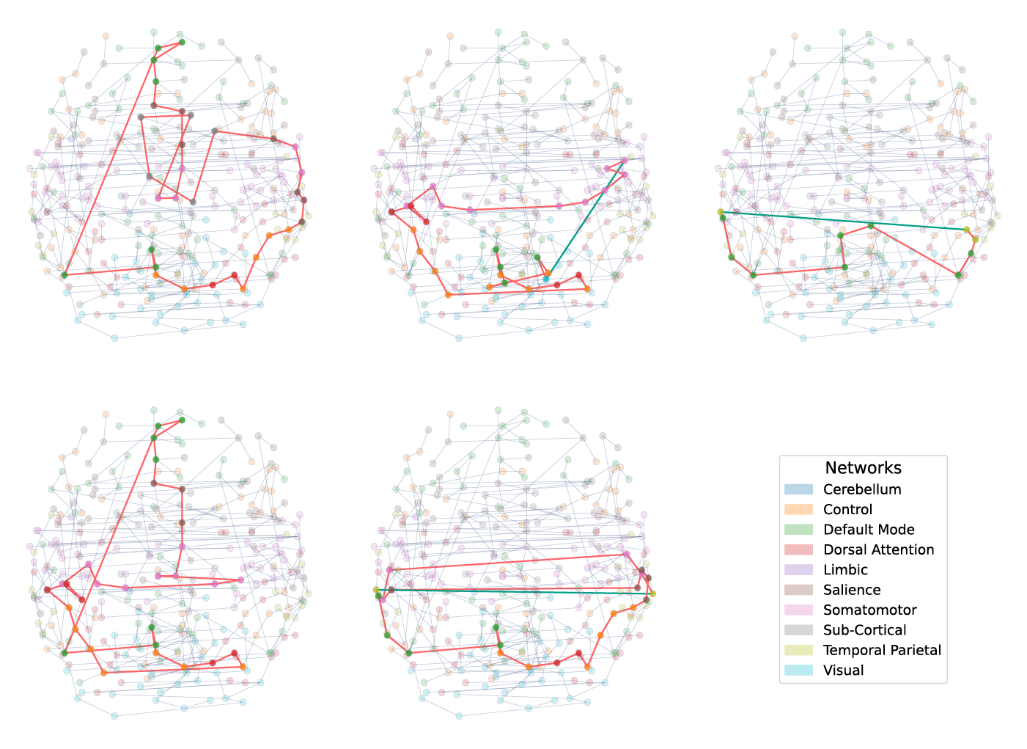

**Figure S10. Visualization of 5 most discriminating 1-cycles in early-onset samples.**

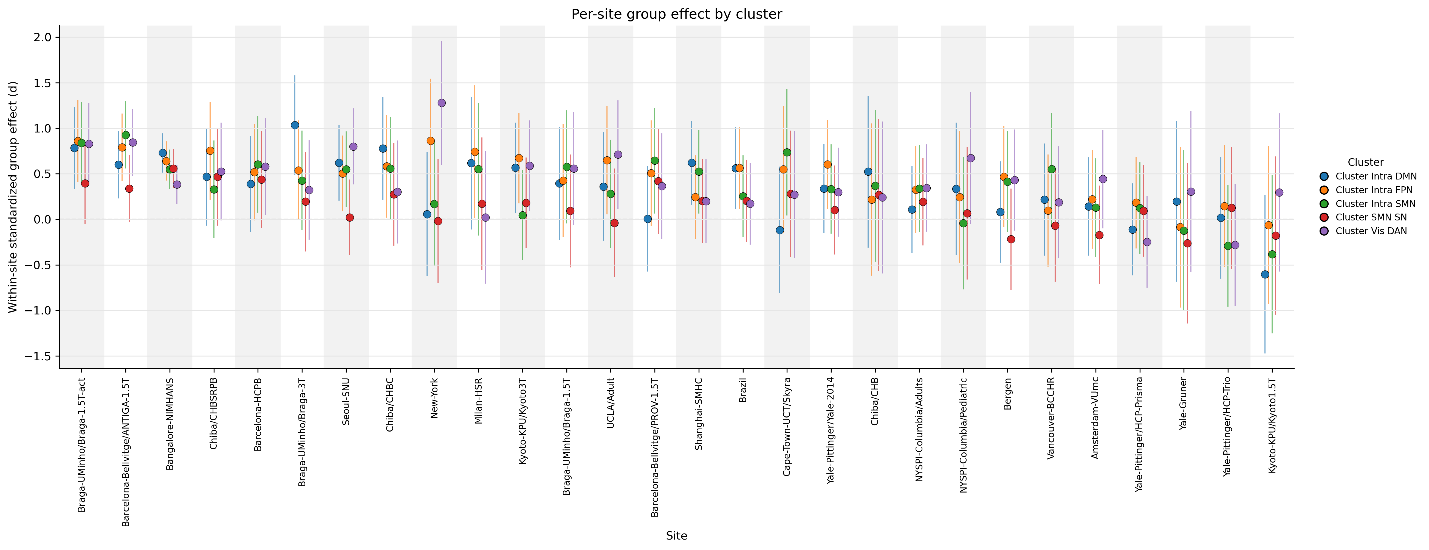

**Figure S11. Within-site OCD–HC group effect for each main-analysis cluster.** For each of the five clusters of significant 1-cycles from the main analysis, the standardized group effect (Cohen's *d*) was estimated separately within every imaging site, using the same general linear model as in the main analysis (diagnosis as the effect of interest; age, sex, and mean framewise displacement as covariates) but without the site term, since each fit is within a single site. Each point is the within-site effect for one cluster (colors), and vertical lines denote the 95% confidence interval; sites are ordered by their mean effect across clusters. Points above the *d* = 0 line indicate the expected OCD-versus-control direction. Across sites the effects were predominantly positive for all five clusters, with wider confidence intervals at smaller sites reflecting lower within-site precision; a small number of sites showed near-zero or reversed estimates, consistent with sampling variability rather than a systematic site effect.

Tables

Table S1(separate file). Available information on sample sizes and scanning acquisition parameters used to obtain structural and functional resting-state data for included ENIGMA-OCD samples.

| Cycle ID | *p*_$\boldsymbol{\alpha}_{\boldsymbol{MST}}$ | *p*_$\boldsymbol{\alpha}_{\boldsymbol{Extra}}$ |
| --- | --- | --- |
| 1821 | 5.38E-03 | 1.94E-02 |
| 1841 | N.S. | 4.53E-04 |
| 6650 | N.S. | 2.11E-06 |
| 7537 | N.S. | 1.32E-03 |
| 11046 | 2.38E-06 | 7.31E-10 |
| 16037 | 2.57E-06 | 2.21E-08 |
| 16885 | 2.85E-08 | 9.14E-07 |
| 21360 | 1.02E-08 | 1.32E-07 |
| 23403 | N.S. | 2.01E-06 |
| 24824 | 2.29E-10 | 5.37E-05 |
| 26488 | 1.12E-05 | 3.30E-06 |
| 29185 | 6.26E-10 | 4.23E-06 |
| 29619 | 2.68E-10 | 1.23E-05 |
| 31045 | N.S. | 3.43E-06 |
| 31972 | N.S. | 6.71E-05 |
| 32280 | 1.49E-02 | 6.22E-04 |
| 32684 | 1.58E-02 | 3.40E-04 |
| 36210 | N.S. | 2.61E-07 |
| 39647 | N.S. | 1.23E-05 |
| 40043 | N.S. | 1.37E-05 |
| 40817 | N.S. | 7.31E-07 |
| 41001 | N.S. | 1.33E-06 |
| 41254 | 5.41E-07 | 1.04E-07 |
| 41316 | 7.43E-05 | 1.22E-06 |
| 41524 | N.S. | 9.13E-10 |
| 41651 | N.S. | 3.81E-06 |
| 41668 | N.S. | 1.83E-04 |
| 41722 | N.S. | 1.12E-04 |
| 41728 | N.S. | 1.32E-06 |
| 41909 | 8.88E-07 | 1.64E-10 |
| 42299 | N.S. | 3.86E-05 |
| 42507 | 1.62E-10 | 2.15E-05 |
| 42837 | N.S. | 4.29E-06 |
| 42846 | 9.50E-05 | 2.00E-07 |
| 42953 | N.S. | 7.72E-05 |
| 43150 | 2.83E-03 | 7.80E-09 |
| 43199 | N.S. | 2.62E-05 |
| 43210 | N.S. | 1.40E-06 |
| 43226 | 3.08E-07 | 1.58E-07 |
| 43617 | N.S. | 9.14E-05 |
| 43959 | 5.17E-03 | 7.33E-07 |
| 44105 | 2.13E-06 | 4.11E-06 |
| 44235 | 2.18E-07 | 1.13E-07 |
| 44375 | N.S. | 2.46E-07 |
| 44572 | 2.35E-02 | 2.97E-08 |
| 44994 | 8.79E-03 | 2.45E-05 |
| 45084 | 9.47E-04 | 1.26E-04 |
| 45127 | N.S. | 2.13E-05 |
| 45489 | N.S. | 1.55E-06 |
| 45653 | N.S. | 1.21E-05 |
| 46052 | N.S. | 7.22E-07 |
| 46054 | 1.01E-07 | 2.20E-07 |
| 46257 | N.S. | 7.94E-07 |
| 46315 | N.S. | 1.93E-04 |
| 46326 | N.S. | 2.32E-06 |
| 46560 | 6.58E-10 | 2.45E-05 |
| 46836 | N.S. | 3.62E-05 |
| 46888 | N.S. | 2.50E-10 |
| 47018 | 5.92E-04 | 9.14E-07 |
| 47061 | N.S. | 2.19E-09 |
| 47219 | N.S. | 6.75E-11 |
| 47279 | 4.53E-08 | 3.44E-05 |
| 47670 | 8.70E-08 | 1.02E-08 |
| 47700 | N.S. | 9.87E-12 |
| 47888 | 6.10E-08 | 4.46E-07 |
| 48185 | N.S. | 5.05E-07 |
| 48190 | 8.63E-07 | 1.00E-08 |
| 48267 | N.S. | 1.66E-08 |
| 48401 | 7.20E-12 | 2.18E-08 |
| 48415 | 4.51E-04 | 4.59E-05 |
| 48484 | N.S. | 3.54E-08 |
| 48491 | N.S. | 2.77E-07 |
| 48591 | N.S. | 3.23E-09 |
| 48817 | N.S. | 6.16E-07 |
| 48825 | 3.92E-07 | 2.61E-07 |
| 48829 | 1.94E-02 | 7.54E-08 |
| 48845 | N.S. | 2.00E-03 |
| 49054 | N.S. | 2.45E-07 |
| 49172 | N.S. | 6.92E-06 |
| 49213 | N.S. | 1.03E-05 |
| 49289 | 9.75E-08 | 3.78E-09 |
| 49302 | N.S. | 2.15E-05 |
| 49321 | N.S. | 1.96E-06 |
| 49469 | N.S. | 2.65E-06 |
| 49580 | 1.12E-05 | 6.36E-07 |
| 49639 | 4.13E-04 | 2.40E-08 |
| 49703 | 3.78E-04 | 2.75E-07 |
| 49849 | 2.76E-02 | 1.19E-08 |
| 49931 | N.S. | 2.66E-06 |
| 49979 | N.S. | 8.43E-07 |
| 49995 | N.S. | 3.77E-06 |
| 50066 | 3.27E-05 | 4.64E-07 |
| 50081 | N.S. | 2.97E-07 |

Table S2. Summary table for *p* values of $\boldsymbol{\alpha}_{\boldsymbol{MST}}$ and $\boldsymbol{\alpha}_{\boldsymbol{Extra}}$ in the most discriminating 1-cycles from the main analysis. All *p* values were false discovery rate (FDR) corrected; significance level was defined by a corrected *p* < 0.05. N.S.: not significant.

| Cluster | Cycles (N) | T | Partial *r* | FDR *q* |
| --- | --- | --- | --- | --- |
| Intra-DMN | 8 | 1.3705 | 0.0439 | 0.2136 |
| Intra-FPN | 27 | 3.1871 | 0.1016 | 0.0074 |
| Intra-SMN | 47 | 1.6429 | 0.0526 | 0.1679 |
| SMN/SN | 1 | 0.0070 | 0.0002 | 0.9945 |
| Vis/DAN | 10 | 1.9987 | 0.0639 | 0.1148 |

**Table S3. Exploratory associations between cluster-level loop expression and Y-BOCS severity in OCD patients.**
